# Systems-level chromosomal parameters represent the suprachromosomal basis for a non-random chromosomal arrangement in human interphase nuclei

**DOI:** 10.1101/052498

**Authors:** Sarosh N. Fatakia, Ishita S. Mehta, Basuthkar J. Rao

## Abstract

Forty-six chromosome territories (CTs) are positioned uniquely in the human interphase nuclei, wherein each of their position can range from the center of the nucleus to its periphery. A non-empirical basis for their non-random arrangement remains unreported. Here, we derive a suprachromosomal basis of that overall arrangement (which we refer to as a CT constellation), and report a hierarchical nature of the same. Using matrix algebra, we unify intrinsic chromosomal parameters (e.g., chromosomal length, gene density, the number of genes per chromosome), to derive an extrinsic effective gene density matrix, the hierarchy of which is dominated by the extrinsic mathematical coupling of HSA19, followed by HSA17 (human chromosome 19 and 17, both preferentially interior CTs) with all CTs. We corroborate predicted constellations and the effective gene density hierarchy with published reports from fluorescent in situ hybridization based microscopy and Hi-C techniques, and delineate analogous hierarchy in disparate vertebrates. Our theory accurately predicts CTs localized to the nuclear interior, which interestingly share conserved synteny with HSA19 and/or HSA17. Finally, the effective gene density hierarchy dictates how inter-CT permutations represent plasticity within constellations and we suggest that a differential mix of coding with noncoding genome may modulate the same.

## Introduction

During interphase stage of cell cycle in human nuclei, chromosomes are confined to a restricted physical location that is referred to as chromosome territory (CT) (Cremer *et al.* 1982; Lanctot *et al.* 2007), which is a small fraction of the entire nuclear volume. Qualitative and quantitative microscopy studies have reported their anisotropic positioning, whose overall format is referred to as the radial arrangement (Cremer and Cremer 2010; Misteli 2010; Bickmore 2013). Interphase CTs not only occupy specific territories in the nucleus, but also intermingle at their borders (Branco and Pombo 2006), while their spatial position remains relatively stable (Zink *et al.* 1998; Tumbar and Belmont 2001; Gerlich *et al.* 2003; Walter *et al.* 2003). In any arrangement of forty-six CTs, which we call a constellation, territories may couple to their neighbors by loci- mediated contacts (Branco and Pombo 2006; Zhao *et al.* 2006) and also via lamin and other nuclear-matrix mediated interactions (van Steensel and Henikoff 2000; Pickersgill *et al.* 2006; Guelen *et al.* 2008). Fluorescent in situ hybridization (FISH) based light microscopy mapping of a clonal population of normal human cells, have reported preferential locations of individual CTs (Croft *et al.* 1999; Sun *et al.* 2000; Boyle *et al.* 2001), and cohorts of two or more CTs (Bridger *et al.* 2000; Cremer *et al.* 2001; Parada *et al.* 2004; Bolzer *et al.* 2005; Mehta *et al.* 2013b). However, significant diversity in preferential positions have been reported for chr5, chr14 and chr21 even in identical cell types (Boyle *et al.* 2001; Cremer *et al.* 2001; Parada *et al.* 2004; Bolzer *et al.* 2005; Mehta *et al.* 2013b), while the preferred positions are markedly different for CTs, such as chr1, chr5, chr11, chr21 and chrY, in diverse cell types (Boyle *et al.* 2001; Cremer *et al.* 2001; Parada *et al.* 2004; Bolzer *et al.* 2005). Hi-C experiments (Lieberman-Aiden *et al.* 2009) and Hi-C-based tethered chromosome capture experiments (Kalhor *et al.* 2012) have reported contact probability distributions of cross linked chromatin and computed a map of inter-CT neighborhoods, which represents the ensemble average of most likely CT constellations. Most importantly, single-cell Hi-C studies have reported variability in the physical structure and spatial conformation of individual CTs (Nagano *et al.* 2013), again using a clonal population, suggesting that the discovery of rare CT constellations (from a clonal population) may be most challenging in a limited population of cells. Therefore, in the context of plasticity within a constellation, a physical basis for nonrandom radial arrangement is essential.

CTs are spatially and temporally regulated and it has been reported that normal chromosomal function significantly impacts their position and vice versa (reviewed in (Bickmore and van Steensel 2013; Cavalli and Misteli 2013; Dekker and Misteli 2015; Pombo and Dillon 2015;Sexton and Cavalli 2015). For example, it has been shown that four distinct CTs undergo statistically significant displacement in the nucleus (relocate from their original position to a new one) after inducing DNA double strand breaks by cisplatin (DNA damaging agent) treatment (Mehta *et al.* 2013b) in an asynchronous population of human dermal fibroblasts. In a significant population of those nuclei, chr17 and chr19 relocated from the interior of the nucleus to the periphery, while chr12 and chr15 were displaced oppositely, from the periphery towards the interior (Mehta *et al.* 2013b). Upon washing off cisplatin from the treated cells original ensemble average arrangement of individual CTs was restored following DNA repair (Mehta *et al.* 2013b). Independent studies, in cancers and genetic instability disorders, have reported that DNA double-strand breaks lead to translocations among closely located CTs (Roix *et al.* 2003). Therefore, a spatially non-random CT constellation directly influence their functions and translocation propensities (Bickmore and Teague 2002). As CT constellations significantly differ in tumor versus normal cells, by virtue of their inter-CT distances (Nikiforova *et al.* 2000; Bickmore and Teague 2002; Parada *et al.* 2002; Roix *et al.* 2003), it is critical to investigate the fundamental basis of the same. An *in vitro* study has been demonstrated that the position of a human chromosome may remain conserved in human-mouse hybrid nuclei and its genes may also be actively transcribed, while sustaining an overall radial form of chromosomal arrangement (Sengupta *et al.* 2008). Therefore, if a species-independent suprachromosomal basis for the non-random anisotropic CT arrangement exists, it must be tenable across disparate vertebrates.

To understand the physical basis of a radial chromosomal arrangement of all forty-six CTs in human nuclei, traditional microscopy-based methods are complemented with *in silico* approaches using polymer-based models (reviewed in (Marti-Renom and Mirny 2011; Fudenberg and Mirny 2012; Halverson *et al.* 2014; Vasquez and Bloom 2014)). It is established that polymers can faithfully represent of chromosomal topology and can reproduce clustered spatial arrangement (Rosa and Everaers 2008; Vettorel *et al.* 2009). In that context, it has been demonstrated that topological constraints govern territorial clustering, because polymers without excluded volume and negligible intermingling have also generated a radial form of arrangement (Dorier and Stasiak 2009; Blackstone *et al.* 2011). Moreover, entropic (nonspecific) factors can lead to anisotropic constellations that may self-organize, as shown using models such as a beads on string model (Cook and Marenduzzo 2009) and a strings and binders model (Barbieri *et al.* 2012). More recently, the role of polymer loops and entropic factors have been used to model and describe CT arrangements in prokaryotes and eukaryotes (Heermann *et al.* 2012; Hofmann and Heermann 2015). These *in silico* methods have used impermeable polymers, and permeable (phantom) ones (Dorier and Stasiak 2009), to derive the theoretical basis of the clustered radial CT arrangement. Based on early investigations (Munkel and Mirny 1998; Munkel *et al.* 1999; Kreth *et al.* 2004) that used intrinsic parameters of the human coding genome, a recent study has modeled radial-like CT clustering in human nuclei (“gene rich” CTs such as chr19 at the nuclear center) by incorporating coding, noncoding and pseudogenes (Ganai *et al.* 2014). More recently, it has been reported that topologically associating domains (TADs) (Dixon *et al.* 2012) are the minimal physical entities of CTs that fold with invariant boundaries across different cell-types (Smith *et al.* 2016), which in turn provide a physical basis for hierarchical folding that encompass domains-within-domains: metaTADs (Fraser *et al.* 2015). Therefore, the physical organization of the human chromosome (or for that matter any eukaryotic chromosome) is known. However, a non-empirical basis for the physical organization, which represents the plasticity and hierarchy of CT constellations, at the suprachromosomal-level has not been reported. Consequently, *in silico* methods have not reported the plethora of CT constellations that exist *in vitro.*

In the context of an interactive chromosomal milieu of the human nuclei, we report a non-empirical suprachromosomal basis for CT constellations that describe their plasticity. We derive systems-level mathematical and physical constraints that are exclusively genome-based, and describe a theory for self-organized CT constellations. Using this theoretical basis for CT constellations, we corroborate their plasticity using inter-CT maps of human nuclei, as obtained using FISH (Croft *et al.* 1999; Sun *et al.* 2000; Boyle *et al.* 2001; Cremer *et al.* 2001; Parada *et al.* 2004; Bolzer *et al.* 2005; Mehta *et al.* 2013b) and Hi-C techniques (Lieberman-Aiden *et al.* 2009; Kalhor *et al.* 2012). As we represent a conglomerate of inter-CT entities, we do not incorporate polymer-based constraints or topology to model the compact non-random radial arrangement of forty-six CTs within the human nucleus. From an evolutionary perspective, we also corroborate the veracity of our predictions, using published FISH results from disparate vertebrates: chimpanzee (Tanabe *et al.* 2002b), mouse (Mayer *et al.* 2005), pig (Foster *et al.* 2012) and chicken (Habermann *et al.* 2001; Tanabe *et al.* 2002a). Using these disparate genomes, we report the theoretical basis for the hierarchy and plasticity within each of their CT constellation, predict the inner CTs in their respective nuclei and subsequently corroborate their identity using previously published results.

## The Theoretical Approach

### Intrinsic parameters of the human genome

The total number of annotated genes per chromosome, and their respective lengths were obtained from National Center of Biotechnology Information (NCBI) Gene database and Mapviewer portal (release 107) for human genome (NCBI 2015). These are parameters are available in Table S4.

### A systems-level representation of intrinsic parameters of the human genome using matrix algebra

For a systems-level theory, we represent the number of genes (coding and noncoding) and the length of chromosomes (traditional intrinsic parameters), for the *N* (= 24) distinct chromosomes of the human genome, as two *N* dimensional column vectors denoted by |***n***⟩, |***L***⟩ and |ℕ⟩, |**∧**⟩ for the *in vitro* and *in vivo* representations respectively. The coupling of these two vectors in the abstract vector space is via a *N × N* matrix. The theory describing the unitary transformation from the *in vitro* to the *in vivo* representation is described at length.

### Unification of intrinsic parameters in the human genome

We denote the different chromosomes of the human genome (chr1, chr2… chr21, chr22, chrX and chrY) as *C_j_* (where 1 *≤j≤ ≤ N =*24). The gene density of *j^th^* chromosome (*c*_*j*_,) is represented by a coupling term *d_j_* in a scalar equation n_j_ = *d_j_L_j_,* where *n*_*j*_ is the total number of genes (that includes protein-coding and noncoding genes) and *L*_*j*_ is the length of *C_j_*. Traditionally, *n_j_* is considered as a parameter that only involves the *j^th^*chromosome, independent of any other *C_k_*, for example, the nearest neighbour of *C_j_*. Therefore, equations involving all intrinsic chromosomal couplings may be collectively represented as 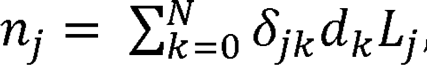 where *δ_jk_* is the Kronecker delta function (=1 for indices *j* = *k,* and =0 for *j* ≠ k). This Kronecker delta formalism enables assignment of null weight to gene density coupling terms for all different chromosomes except *C*_*j*_. As we shall show later, the formalism provides us with the scope for the intrinsic parameters of to be *C*_*j*_ influenced by another *C*_*k*_, thereby providing systemic-level constraints, which is otherwise not apparent. Using this mathematics, we formulate a theory that couples intrinsic parameters of different chromosomes to derive their systems-level crosstalk. For now, to facilitate this systems-level approach in an *in vitro* setting, we posit 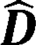 as a diagonal *N × N* matrix with intrinsic gene density (*d*_*j*_) as the diagonal elements (real eigenvalues of 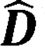). It must be noted that intrinsic parameters are experimentally obtained from cytogenetic as well as sequencing efforts under laboratory conditions and they represent an *in vitro* milieu. Therefore, these parameters may not accurately represent typical systems biology of the *in vivo* milieu that nuclei manifest. For brevity, we denote the set of basis vectors for the *in vitro* or diagonal representation as 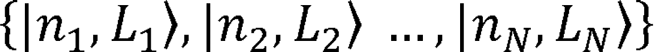, labeled by the intrinsic parameters: total gene count per chromosome (*n*), its length (*L*) in megabase pair (Mbp) and a subscript *j* for referencing *C_j_*. (Intrinsic chromosomal parameters are provided in Table S4). A systems-level formalism in a dimensional vector space using a *in vitro* coupling matrix is presented as a matrix equation: where,
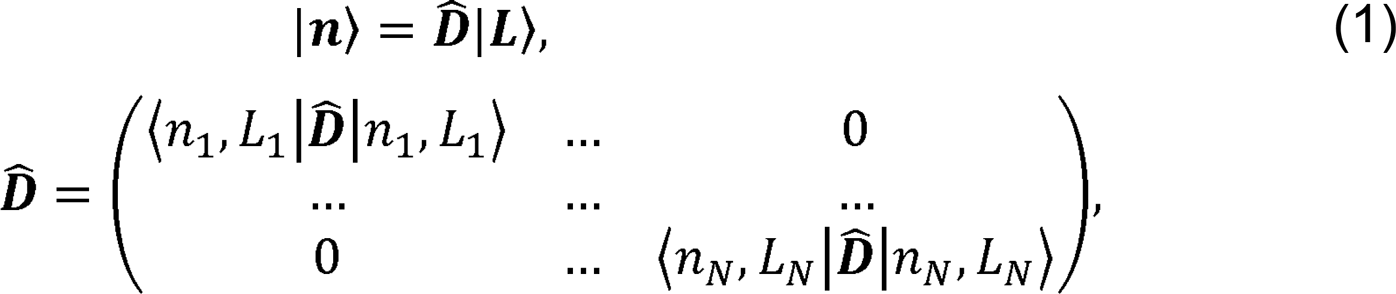

such that the intrinsic gene density 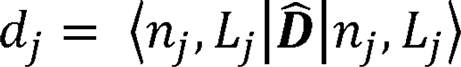 (megabase pair Mbp^−1^ units). Here 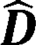 is a diagonal matrix and ❘***n***⟩ and ❘***L***⟩ are *N* × 1 matrices (or column vectors) whose components represent the *in vitro* intrinsic parameters, namely the number of coding genes and chromosome length. As the *in vitro* gene density space does not describe inter-chromosomal couplings, the hierarchical nature of extrinsic gene density remains implicit.

### Extrinsic parameters of the human genome

Now, we mathematically describe extrinsic chromosomal parameters in an *in vivo* context. We used the intrinsic parameters from all the unique *N* (= 24) human chromosomes (22 autosomes and X, Y chromosomes) to quantify the crosstalk among the 46 CTs (diploid genome) within the human nucleus. Systems-level coupling among them may be represented in a *N* dimensional *in vivo* vector space defined by a *N × N* Hermitian matrix 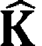. This space is defined by a set of *N* basis vectors (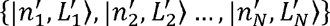) labeled by extrinsic systems-level parameters: effective gene (*n*′) count and effective length (*L*′) in Mbp, along with subscripts that denote labels from column and row vectors. Next, analogous to the *in vitro* model (Equation 1), we represent an abstract effective gene count and effective length as vectors denoted by |ℕ⟩ and |**∧**⟩ respectively. We posit:

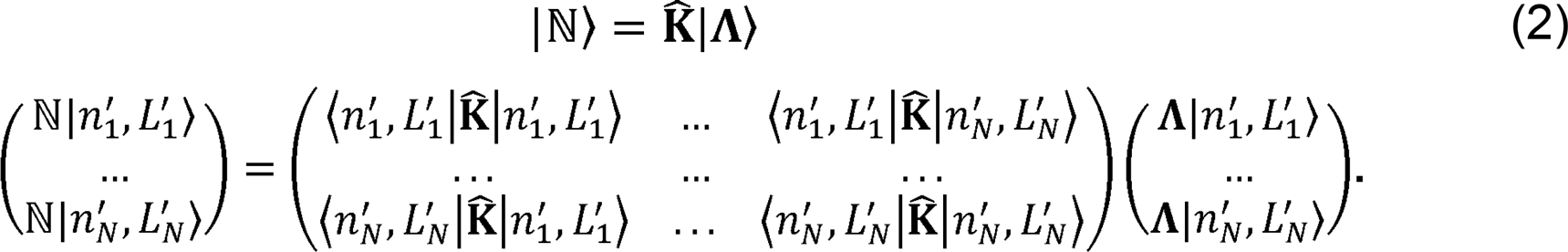

We hypothesize that the systems-level coupling of CTs is commutative and associative, such that it may be represented by a matrix (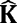), which is a real, symmetric Hermitian matrix. We hypothesize that this coupling matrix is due to the nearest-neighbour inter-CT coupling of *C*_*j*_) with *C_k_,* hence in Equation 2 the original generic coupling matrix 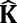 may be approximated to the first-order, using 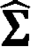, where 
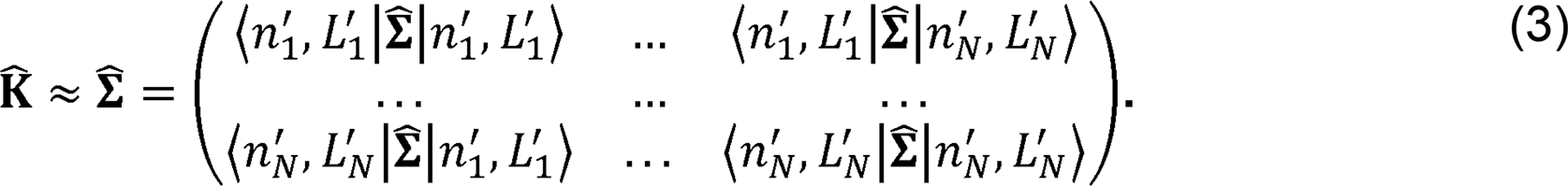

The elements of 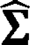 are the extrinsic coupling coefficients, and physically they virtually represent the spatial inter-CT crosstalk. Next, we derive these extrinsic couplings using intrinsic chromosomal parameters: number of genes (*n*_*j*_), chromosomal length (*L*_*j*_), gene density (*d*_*j*_), along with *N* scalar equations *n*_*j*_ = ∑_*k*_δ_*jk*_*d*_*k*_*L*_*j*_, with *d*_*j*_>0,*L*_*j*_ >0, where *δ_jk_* is the Kronecker delta function. To explicitly derive the obscure inter-CT “mixing” (which we refer to as biological crosstalk), we represent 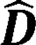 in the *in vivo* vector space. This deconstruction is obtained, using the Spectral Theorem, as a symmetric *N* × *N* matrix 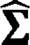 such that: 
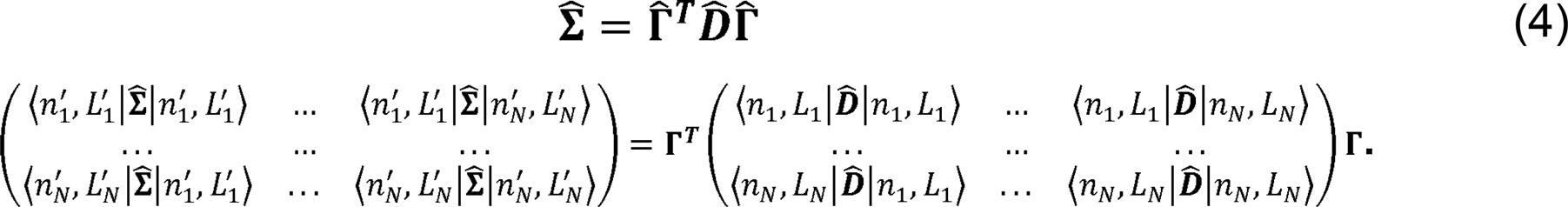

Here, the *j* row and *k* column element of 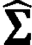 is denoted as 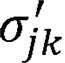, in megabase pair (Mbp) units, (where 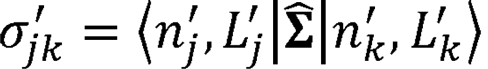), and 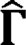 is a *N* × *N* orthogonal matrix ( 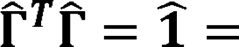 unitary matrix).

### Hierarchical clustering of effective gene density

To deconstruct the hierarchy of effective gene density matrix, we used five different hierarchical clustering algorithms implemented in R software-package (hclust function that is based on Fortran code contributed to STATLIB by Murtagh). The different algorithms used were (A) complete linkage, (B) single linkage, (C) centroid linkage, (D) average linkage and (E) Mcquity implementation.

## Results

The traditional schematic of the human genome is the linear DNA sequence of nucleotides, which includes the twenty-four different chromosomes in sequential order (chr1, chr2, …, chr21, chr22, chrX, and chrY). In this report, we consolidate the intrinsic parameters of all chromosomes in an abstract D vector space and mathematically derive extrinsic inter-CT parameters (representative of inter-CT biological crosstalk or the mathematical coupling among different CTs). This abstract formalism is used to identify systemic extrinsic constraints among the forty-six CTs in an interactive milieu, which constitute a CT constellation that is non-random. Moreover, we also show that the inter-CT coupling supports an identical hierarchical arrangement within every unique constellation, which arises along with permissible degeneracy (ambiguity due to CT permutations), as in a classical statistical ensemble of accessible CT constellations. This report builds upon our original work (Fatakia *et al.* 2015) and extends the formalism to other disparate eukaryotic species.

### Systems-level inter-CT coupling coefficient represents effective gene density

Here, we report the mathematical formalism to depict all unique inter-CT pairs in an *in vivo* context, which is distinct from the *in vitro* context, where individual CTs are accessed without the influence of other CTs (Materials and Method). For notational brevity for *in vitro* representation, we denote the different chromosomes of the human genome (chr1, chr2… chr21, chr22, chrX and chrY) as *C_j_* (where 1 *≤j ≤ N =*24). In theory, any CT may pair with any other CT as in a CT constellation, so we derive all extrinsic inter-CT parameters that characterize constellations in an *in vivo* representation. We hypothesize that for each nearest-neighbour pair 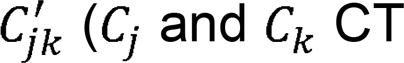(*C_j_* and *C_k_* CT pair) the composite effective gene count (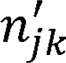) or paired chromosome’s gene count(PCGC) is mathematically coupled to an effective length 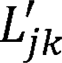 via an effective genome gene density 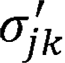 as: 
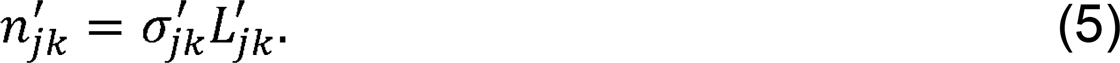

We define the dimensionless PCGC parameter 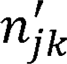 as the harmonic mean of the total number of annotated genes, from coding and noncoding parts, in 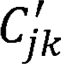. The harmonic mean (as opposed to geometric mean or arithmetic mean) is an ideal statistic that represents very diverse number of genes per chromosome, as it gives lower weightage to very high values. Therefore, the effective gene count for a given 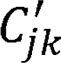 is represented in terms of its intrinsic parameters *n_j_* and *n_k_* as: 
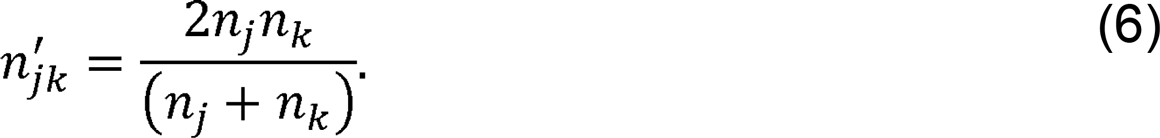

Similarly, to best represent the size or length of 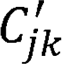, we define the effective length 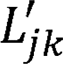 as the harmonic mean of intrinsic parameters *L_j_* and *L_K_*:

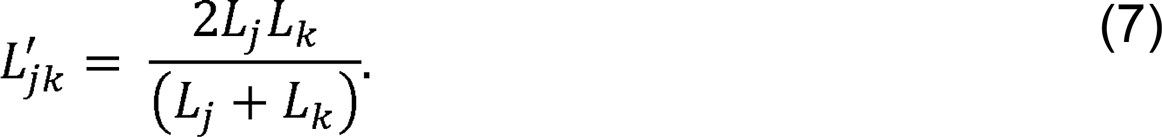

Representing 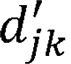 as a harmonic mean of intrinsic average gene densities:

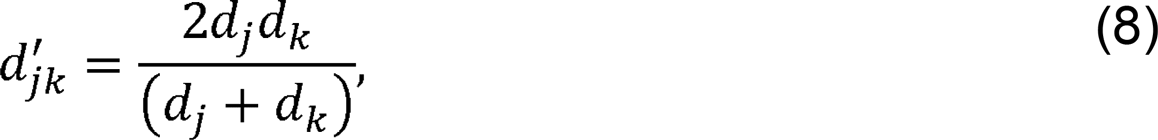
and using Equations 5 – 8, we derive the extrinsic inter-CT coupling coefficient as:

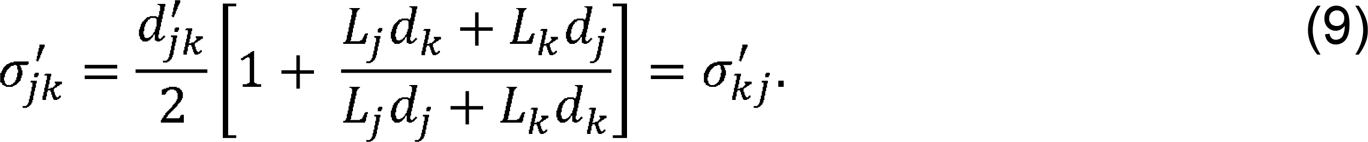

This inter-CT spatial coupling coefficient parameter 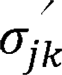 has the physical dimensions of gene density and is henceforth referred to as the effective gene density. It represents the extrinsic coupling within paired nearest-neighbor 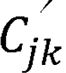 an inter-CT *in vivo* unit with an effective length (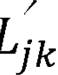). To access the contributions of all inter-CT pairs in a genome, we normalize 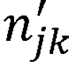 in Equation 5 with two genome-specific normalization constants: *M*( = 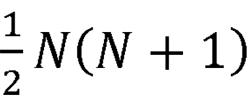 and 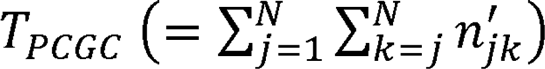. When expressed as a percentage the normalized Equation 5 is:

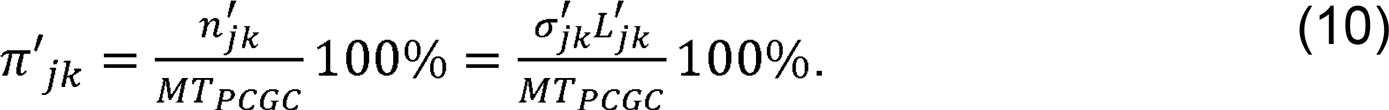

### Sum total of all the extrinsic effective gene density parameters in human genome greatly exceed intrinsic ones

Our minimalistic iVdimensional (24D) vector model invokes abstract unified entities 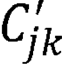 instead of solitary CTs. In Equation 9, the derived effective gene density for 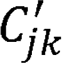 has the conventional term (*L_j_d_j_*) and an unconventional “mixed” term *(L_j_d_k_),* representing a systems-level coupling of CTs. All effective gene density (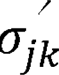) parameters are represented using the effective gene density matrix – a *N × N* real symmetric matrix 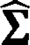 (Materials and Methods), with 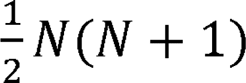 unique elements. Each of those elements (parameters) is derived using the genomic information. Of those 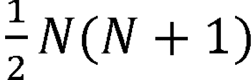 unique parameters, *N* are intrinsic (diagonal elements) and 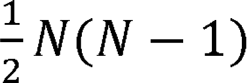 are extrinsic (off-diagonal elements). Hence, in human nucleus, where *N* = 24, the maximum possible number of extrinsic systems-level parameters is nearly an order of magnitude grater (twelve times) than that from the more familiar *in vitro* model. The theory presented here involves the couplings among homologous CTs (modeled by conventional term) and non-homologous pairs (modeled by “mixed” term). Therefore, this matrix method models the biological crosstalk, and as the sum total of extrinsic parameters far exceed the intrinsic ones, we have identified a larger number of mathematically derived genomic parameters that pair up (mathematically couple) chromosomal length with gene density for homologous and non-homologous chromosome partners within the nucleus.

### Effective gene density matrix constrains human CT arrangement

We represent extrinsic inter-CT coupling coefficient, 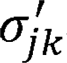, as off-diagonal (*j,k*) elements of the systems-level coupling matrix (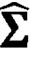) (Materials and Methods). Using Equations 8 and 9, each matrix element 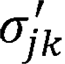, for 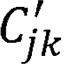, is:

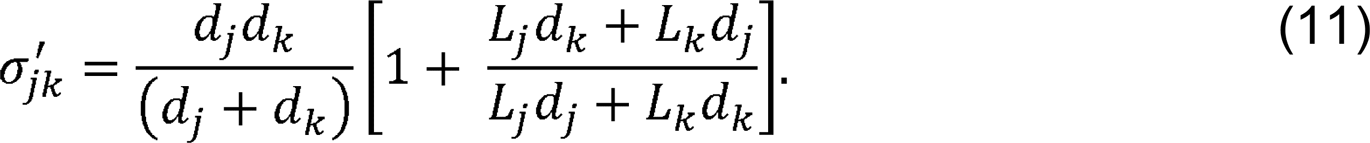

The large diversity in effective gene density matrix is represented as a histogram (Figure 1A) and hierarchically clustered heatmap (Figure 1B). The histogram for effective gene density is asymmetric, skewed toward higher values. The highest effective gene density for 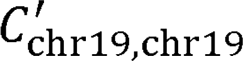 is statistically significant from the histogram mean (its difference from the mean value is 4.7 times the root-mean-square deviation of the histogram). Effective gene density heat map substantiates that for a given CT; there may be variability in its extrinsic gene density that is contingent on the neighborhood of CTs (left-most column and lower-most row in Figure 1B). For example, effective gene density couplings for 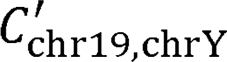 are much lower (less than fifty percent) than the effective gene density couplings for 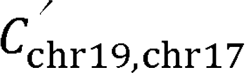. This implies that if extrinsic (effective) gene density constraints were to dictate the preferential positioning of CTs, then the co-occurrence of chr17 and chr19 as pair would be strongly biased, and therefore more likely, in contrast to the chr19 and chrY inter-CT pair.

**Figure 1.**
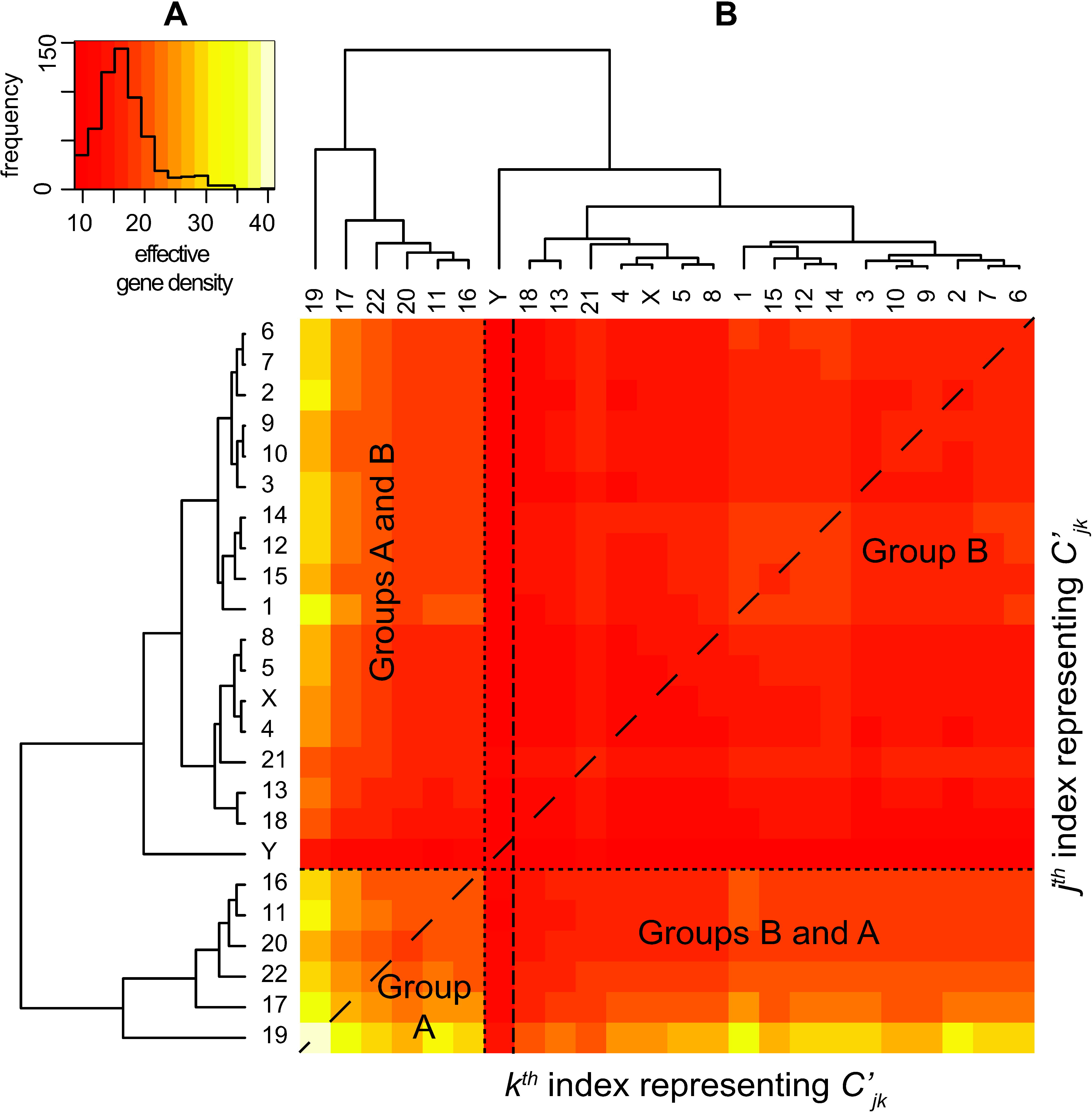
Effective gene density obtained from extrinsic inter-CT pairs of the human genome. A histogram of effective gene density from all unique chromosome pairs 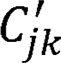 is represented in **A**. The color key for the histogram (red to yellow to off-white) corresponds to increasing effective gene density values on the x-axis. A hierarchically clustered heatmap of effective gene density matrix from all 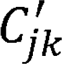 chromosome pairs, and is indexed by labels and on the y- and x-axes respectively, is shown in **B**. Values of effective gene density from 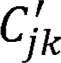 and 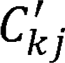 are identical, indicated by a dashed diagonal line, and the horizontal and vertical dashed lines segregate inter-CT pairs where both *C_j_* and *C_k_* are from Group A, with those from Group B, or their admixture. Here, HSAY (human chrY), a Group B CT is highlighted by two vertical dashed lines, which represent its unique position in the overall hierarchy. The color key for effective gene density values in panels **A** and **B** are consistent.

Using the genome-specific normalized PCGC values (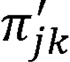); we wanted to determine if neighborhood CT effects could be used to delineate spatial CT arrangements. The *in vitro* spatial positions of individual CTs were already known from a high volume microscopy study performed using dermal fibroblast nuclei (Mehta *et al.* 2013b). Hence, we sought to identify patterns of hypothetical extrinsic couplings among all inter-CTs. Using the microscopy-based data from fibroblast nuclei (Table S1), we computed 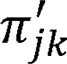 versus 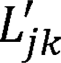 (effective length) for cases in which *C*_j_ and *C*_k_ were both exclusive to (i) the nuclear interior, (ii) the periphery, and (iii) spatially intermediate to (i) and (ii). Such a scatter-plot of 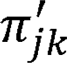 versus 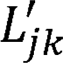 (Figure 2A), whose composition is fibroblast cell-type specific, confirms the segregation among paired CTs that are in the inner core versus the periphery of the nucleus, suggesting a hierarchy in PCGG versus effective length for 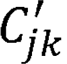. The scatter-plot retrieved for fibroblast also describes an intermediate category, which depends on whether C_j_and *C_k_* is interior or peripheral. However, it is important to note that the cluster of interior versus periphery is different in lymphocytes (Table S1), leading to commensurate changes in the intermediate category (Figure 2B). Interestingly, 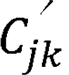 that do not overlap with instances where both CTs are confined to the interior / intermediate / periphery of the nucleus are also represented in the scatter-plot. A CT from any one of the above three spatial zones coupled with another CT from a different zone constitutes the non-overlapping category (Figure 2). Therefore, we hypothesize that a systematic hierarchy and degeneracy in spatial CT arrangement is obtained when inter-chromosomal coupling via effective gene density is computed. We surmise, while the scatter-plot in Figure 2 is representative of the overall allowable pattern, the cell-type specificity dictates the details of CT positioning between the nuclear interior versus intermediate versus peripheral zones.

**Figure 2.**
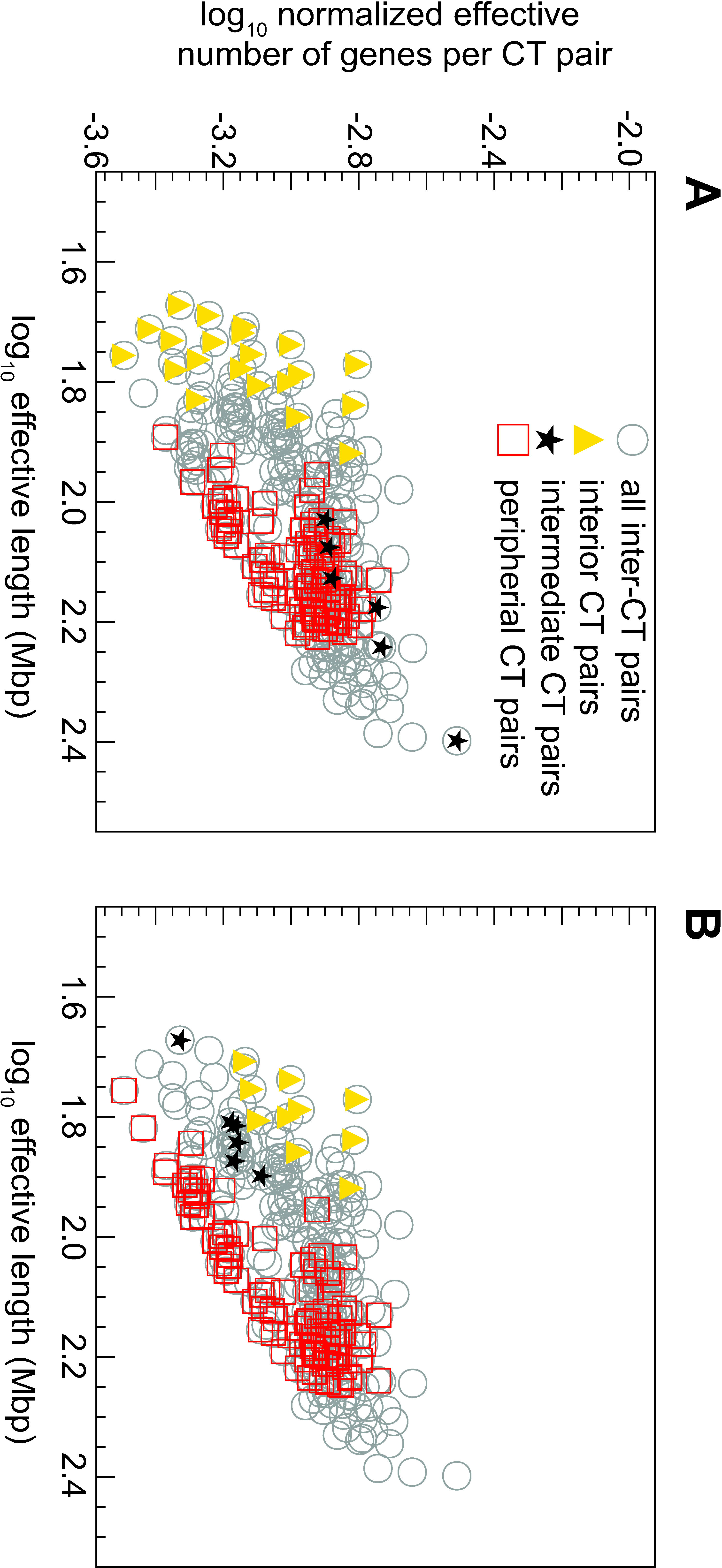
Normalized effective number of genes versus effective length for inter-CT pairs. The graph of normalized effective number of genes (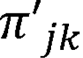) for every 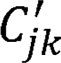 pair versus their effective length 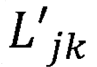 for all 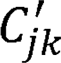 is shown as a circle (open circle). Panel **A** represents the preferential CT location data from fibroblast nuclei and **B** represents lymphocyte data. The coordinates representing 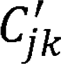 pairs, wherein both CTs are from nuclear interior (triangle), periphery (box), and spatially intermediate region (star) are superimposed over those representing all pairs (open circle). The ones shown in open circle that do not overlap with either a triangle, box, or star, represent those pairs where CTs belong to different mix of spatial categories. Only unambiguous spatial CT positions are represented using Table S1.

### Hierarchy of effective gene density from the human genome supports CT constellations

As normalized PCGC (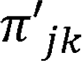) and effective length (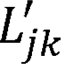) terms are coupled via the effective gene density 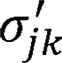 (Equation 10), we investigate its hierarchy in the context of CT arrangements in human interphase nuclei. We collate pairs of CTs with similar average effective gene density to compute a binary tree in effective gene density space. As effective gene density is a mathematical function of the CT length and gene density, such a binary tree hierarchically clusters CTs based on their extrinsic systems-level attributes. Hence, we used the complete-linkage algorithm (Materials and Method) to compute the hierarchical clustering within the abstract effective gene density space. Our analysis of the human genome led to a segregation of effective gene density from CTs into two primary clusters, Groups A and B (Figure 1B, Figure 3). We represent Group A with chr11, chr16, chr17, chr19, chr20, chr22 and Group B with the remaining 18 CTs (Figure 3).

**Figure 3.**
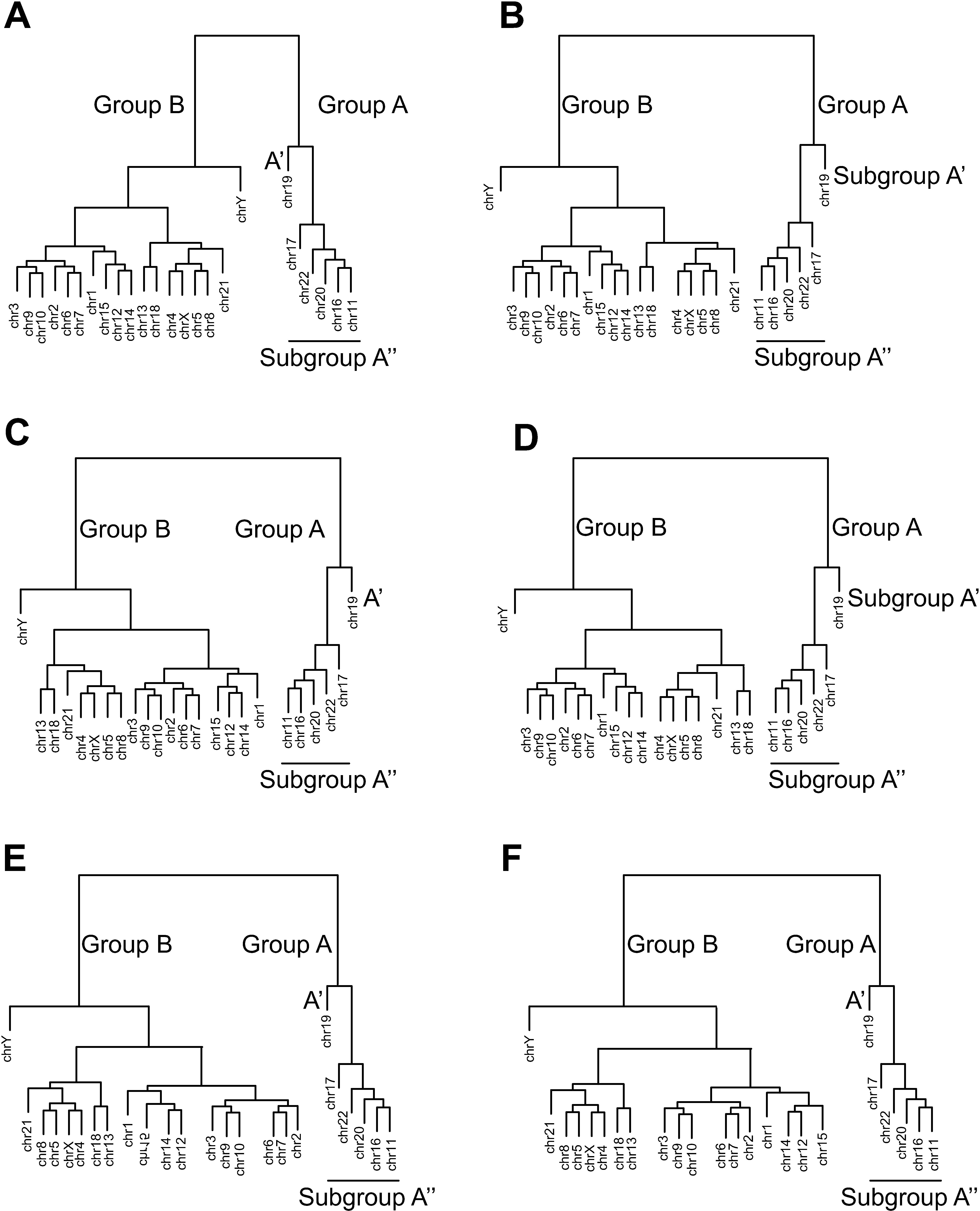
Equivalent hierarchically clustered dendrograms derived from effective gene density matrix. The inter-CT coupling dendrograms derived from effective gene density matrix for *Homo sapiens* (using coding and noncoding genome) represent unique CT constellations in **A-F**. These six unique CT constellations can be perceived on an abstract 2D plane by consigning HSA19 (human chr19) to the nuclear interior, as in **A-F**. The vertical lines connote hierarchy in evolutionary distance or evolutionary diversity suggesting CTs with diverse effective gene density are evolutionary distant. The length of horizontal branch does not represent any hierarchical information. Interior and peripheral CTs are denoted as Groups A and B respectively. Although chr21 and chrY are Group B CTs and relatively gene poor, the theory supports a constellation with them as interior chromosomes due to neighbourhood CT effects (**A**). A hierarchical and degenerate representation enables the rationalization of chr21 and chrY as interior CTs (A) such as in fibroblasts versus peripheral CTs such as in lymphocytes (**B**). Similarly, chr11, chr1, and chr14 are rationalized as intermediate to peripheral CTs (**B** versus **C**). Constellations with chr18, and chr2 towards the nuclear interior are illustrated in (**D**) and (**E**) respectively. Although, chr2 and chr18 are primarily located at the nuclear periphery, (**D**) and (**E**) are instances when they are located differently. The constellation (**F**) illustrates a sub-cluster of chr12, chr14 and chr15 in the neighbourhood of Group A CTs.

### HSA19 dominates hierarchy of inter-CT effective gene density and dictates spatial CT position in human interphase nuclei

The inter-CT effective gene density of HSA19 (chr19) with every other CT represents the entire range of permissible effective gene density values (Figure 1A). Therefore, chr19 represents the primary hierarchy of all inter-CT effective gene density parameters. Interestingly, if we consider a hypothetical pruning of chr19 (from Group A), then the hierarchy of Group A changes completely with respect to Group B. To avoid any algorithmic bias, which may be introduced as an artifact of the clustering method, we recomputed the effective gene density matrix hierarchy using single-linkage (Figure S1B), centroid-linkage (Figure S1C), average-linkage (Figure S1D), and Mcquity (Figure S1E) algorithms. We report chr19 consistently maintains the primary hierarchy as obtained from each of these different algorithms. As chr19 occupies a unique and primary hierarchy among all 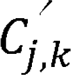, one could represent it at the origin of an abstract effective gene density space, or consign it to the interior of the nuclear volume in the context of a particular CT constellation. Once the physical location of the CT(s) with primary hierarchy is known within the nucleus, our mathematical formulation describes the subsequent hierarchical basis for self-organization of the other CTs with respect to that. It has already been established that chr19 is at the interior of the human nucleus using fibroblasts (Croft *et al.* 1999; Sun *et al.* 2000; Boyle *et al.* 2001; Cremer *et al.* 2001; Bolzer *et al.* 2005; Mehta *et al.* 2013b), lymphocytes (Croft *et al.* 1999; Boyle *et al.* 2001; Cremer *et al.* 2001; Bolzer *et al.* 2005) and lymphoblastoids (Lieberman-Aiden *et al.* 2009; Kalhor *et al.* 2012) cells. Therefore, this localized one-to-one mapping of CTs in an abstract effective gene density space with the real spatial nuclear volume facilitated subsequent spatial arrangement of the other CTs with respect to each other as well as with chr19, such that a CT constellation may be conceived (from the interior towards the periphery). Our results imply that the primary hierarchy of chr19 in effective gene density space is essential for the overall non-random and hierarchical spatial arrangement of other CTs in the human nucleus. Therefore, Group A was subdivided into Subgroup A’ (chr19) and Subgroup A’’ (chr11, chr16, chr17, chr20 and chr22) (Figure 3).

We report that the hierarchy of Subgroup A’’ was juxtaposed intermediate to chr19 and Group B (Figure 3), which implies that CTs of Subgroup A’’ (chr11, chr16, chr17, chr20 and chr22) were constrained to be adjacent to Subgroup A’ (chr19) in a non-random anisotropic constellation. In addition, the hierarchical clustering supports degenerate representations of CTs 11 and chr16 in the neighborhood of other CTs (Figure 3). The remaining eighteen CTs (Group B), which constituted a dominant fraction, were segregated from Group A’, except for chrY. The dendrogram leaf representing chrY stands out juxtaposed between the six Group A CTs and seventeen Group B CTs (Figure 3). Noticeably, on the heat map of Figure 1B, diversity in effective gene density among 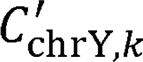 pairs are relatively lower (low variability) compared to corresponding results from 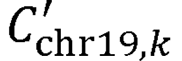 (greater variability). Clearly, in contrast with that of chr19, if chrY were pruned from the dendrogram, the hierarchy of Group A with respect to Group B does not alter (Figure 3). These results support three important consequences of CT arrangements in human nuclei. Firstly, it demonstrates that our theoretical basis for suprachromosomal CT constellation is identical in diploid 46,XX female nuclei (Figure 4) and 46,XY male nuclei (Figure 1). Secondly, contingent on near-neighbor effective gene density couplings, the cell-type, and the poised state of genes on CTs, we predict that the spatial position of chrY with respect to chr19 can lead to great variability: they can be interior CTs as in fibroblasts (Figure 3A) (Mehta *et al.* 2013b), or distantly placed toward the periphery as in lymphocytes (Figure 3B–F) (Boyle *et al.* 2001). Thirdly, and most importantly, the “orphan” nature of chrY, for example, with no homologous partner for crossing over (except with the pseudo-autosomal region in chrX), and its functional specialization for spermatogenesis is also consistent with our systems-level effective gene density hierarchy analyses. The spatially “delocalized” nature of chrY predicted here is indicative of special evolutionary trajectory of *Homo sapiens* chrY (HSAY), which is very distinct from chrY in *Pan troglodytes* (PTRY) (Hughes *et al.* 2010; Cortez *et al.* 2014).

**Figure 4.**
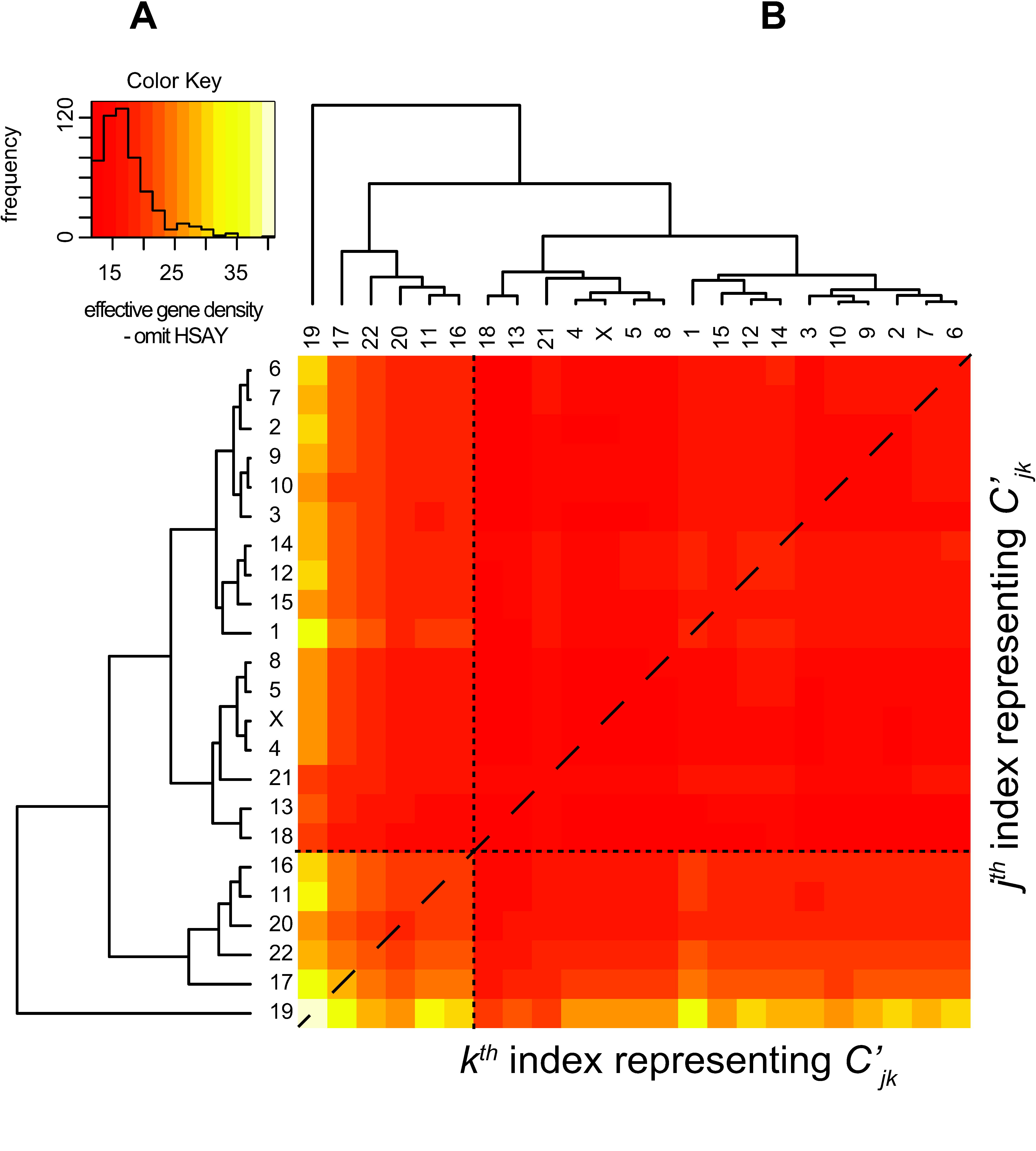
Effective gene density matrix obtained from the (46,XX) diploid female genome. The hierarchy of extrinsic inter-CT effective gene density matrix obtained using the intrinsic parameters of all the twenty-three unique chromosomes that represent the 46,XX human female genome. Inset histograms for these panels represent effective gene density color key.

### Corroboration of theoretical results

Next, we corroborate all our theoretical results in three independent ways. First, these results are corroborated using FISH-based microscopy results of reported CT arrangements in human interphase nuclei (Croft *et al.* 1999; Bridger *et al.* 2000; Sun *et al.* 2000; Boyle *et al.* 2001; Cremer *et al.* 2001; Parada *et al.* 2004; Bolzer *et al.* 2005; Mehta *et al.* 2013b).

### (i) Corroborating plasticity of CT constellations using FISH results

There is plasticity (ambiguity in preferential positioning) in CT position of chr1, chr14 and chr21 from FISH studies (Boyle *et al.* 2001; Cremer *et al.* 2001; Bolzer *et al.* 2005; Mehta *et al.* 2013b), even within a clonal population. Using published reports (Boyle *et al.* 2001; Mehta *et al.* 2013b), we have compiled the inherent variability in preferential CT locations for nine CTs: chr1, chr5, chr6, chr8, chr15, chr16, chr20, chr21 and chrY, in lymphocytes and fibroblasts (Table 1). A physical basis for ambiguity and variability is rationalized here using our PCGC model. We delineated seventeen CTs from Group B (excluding chrY) as largely equidistant *en masse* from Group A, (primarily in the context of chr19) in effective gene density space. However, due to degenerate configurations, it is plausible that cohorts of CTs in Groups A and B will have an effective gene density-based altered spatial arrangement (for example with respect to chr19) and still maintain effective gene density hierarchy (Figure 3). Such a hierarchical and degenerate representation has enabled ambiguous yet non-random chromosomal spatial arrangement, which has been experimentally discerned but not rationalized so far. As shown earlier in Figure 3A, the position of chrY in the dendrogram facilitates its spatial locations in the neighborhood of six CTs from Group A (largely gene rich CTs) as well as the other seventeen CTs from Group B (largely gene poor CTs) as represented in the two equivalent dendrogram representations. Similarly, it can also be argued that near-neighbor effects will give rise to dissimilar CT neighborhoods or constellations that represent chr21 in effective gene density space (Figure 3A and 3C), the outcome of which can give rise to contrasting spatial positions (interior versus periphery) in fibroblasts and lymphocytes (Boyle *et al.* 2001; Mehta *et al.* 2013b) (Figure 2).

**Table 1.**
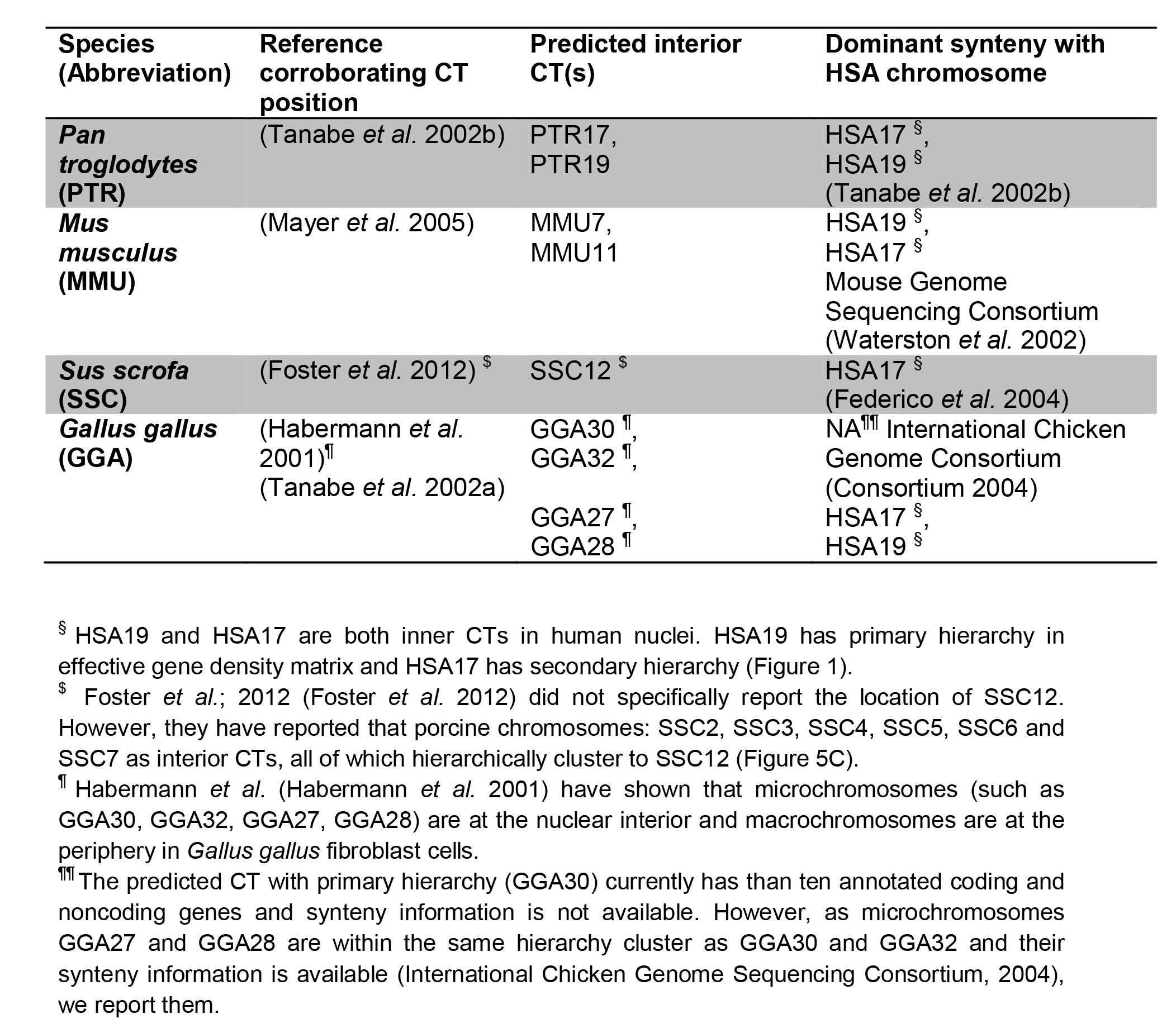
Interior CTs predicted using hierarchical clustering of effective gene density matrix. CT(s) with primary hierarchy in effective gene density are located in the nuclear interior and influence CT neighborhoods forming a non-random CT constellation. HSA, PTR, MMU, SSC and GGA are standard identifiers used to represent the human *(Homo sapiens),* chimpanzee *(Pan troglodytes),* mouse *(Mus musculus),* pig *(Sus scrofa),* and chicken *(Gallus gallus)* chromosomes respectively.

Consider another CT constellation that includes chr11 (Figure 3B), where we can justify the formalism for neighborhood CT effects by positioning it with chr1 and chr14 as intermediate to peripheral (Figure 3C). For example, chr11 in the vicinity of chr1 or chr14 (Figure 3C) has lower effective gene density, as opposed to chr11 in the vicinity of chr16 or chr20 (Figure 3A) as CTs couple differently among themselves in these two scenarios (a lower effective gene density neighborhood occurs if the gene density disparity among them is large and vice versa). Using the same physical basis that justifies various CT constellations, we can represent chr18 or chr2 towards the nuclear interior (Figure 3D, 3E) and likewise the sub-cluster of chr12, chr14 and chr15 (Figure 3F). Clearly spatial positioning of CTs can be very different, particularly when cell- specific instances and poised state of genes are considered. Permutations among CTs with degenerate effective gene density neighborhoods may have negligible consequence on a pan-nuclear scale. Hence, such degenerate CT permutations may be more feasible as opposed to those involving non-degenerate effective gene density CT neighborhoods. Therefore, our theory provides the physical basis for diverse anisotropic constellations of CTs at the suprachromosomal level.

### (ii) Corroborating hierarchy of effective gene density using Hi-C results

Lieberman-Aiden *et al.* have computed inter-CT contact probabilities for normal human cell lines (Lieberman-Aiden *et al.* 2009). Specifically they have reported that the small gene-rich CTs: chr16, chr17, chr19, chr20, chr21 and chr22 preferentially interact with each other. From this group except for chr21, all other CTs match up with our results (Table S2). This cluster is represented in our analysis as Group A (Figure 1 and Figure 3). We rationalize that near-neighbor effects in our PCGC theory, as illustrated in Figure 3A, may lead to the omission of chr21 from the Group A and consigned to Group B in our analysis. In that scenario, chr21 is placed at the edge of Group B CTs, adjacent to Group A. In addition, our Group A also contained chr11, which was not supported by the Hi-C result. Interestingly, in the tethered chromosome capture (TCC) technique, which is a derivative of Hi-C experiment that sought to improve the signal to noise ratio, Kalhor *et al.* (Kalhor *et al.* 2012) have showed chr11 is part of an inner cluster of CTs, consistent with our Group A CTs (Table S2). So it is justifiable that the effects of neighborhood-CTs and cell-type specificity constellations, and the sensitivity of experimental method dictate how well we resolve the unique from rare constellations. In that report, inter-CTs contact probabilities were used to compute two clusters, which they referred to as Cluster 1 and Cluster 2 (Kalhor *et al.* 2012). The reported Cluster 1 contained 10 CTs, which included chr11, chr16, chr17, chr19, chr20 and chr22 - all of which are Group A CTs. However, in addition chr1, chr14, chr15 and chr21 were also part of Cluster 1. In fact, chr1 (the largest chromosome in the human genome) is represented as one that may be localized proximal to Group A CTs in our hierarchy analyses (Figure 3C), a result consistent with chr1 being part of inner core CTs (Kalhor *et al.* 2012). Therefore, we conclude most of our results of constellation plasticity and the hierarchical clustering of CTs have also been supported by these unbiased Hi-C experiments.

### (iii) Corroborating interior CTs using disparate mammalian genomes

Next, we used four different model organisms, which have been shown to exhibit radial-like CT arrangement, to compute primary hierarchy of CT(s) in the abstract effective gene density space. Using PCGC analysis, we predicted interior CTs in *Pan troglodytes* (Figure 5A), *Mus musculus* (Figure 5B), *Sus scrofa* (Figure 5C) and *Gallus gallus* (Figure 5D) nuclei. We corroborated our predictions using previously reportedly FISH results for chimpanzee (Tanabe *et al.* 2002b), mouse (Mayer *et al.* 2005), pig (Foster *et al.* 2012) and chicken (Habermann *et al.* 2001; Tanabe *et al.* 2002a). We apply our theory to these vertebrates as their radial CT arrangement has been previously studied, and predict their inner core CTs. Without using any sequence information, we report that PTR17; PTR19 (in chimpanzee), MMU7; MMU11 (in mouse), SSC12 (in pig), and GGA30, GGA32, GGA27, GGA28 (in chicken) have primary hierarchy. All of these CTs have been identified as interior CTs (Tanabe *et al.* 2002b; Mayer *et al.* 2005) (Table 1). However, as shown in Table 1, for two species the spatial position of CTs with primary hierarchy have not been mapped using FISH technique, but we report that they hierarchically cluster to other CTs that have been mapped as interior CTs: pig chromosome SSC12 (Foster *et al.* 2012) and chicken microchromosomes GGA30, GGA32 (Habermann *et al.* 2001; Tanabe *et al.* 2002a). Next, one can identify the human chromosomes, which have a dominant synteny with predicted inner CTs to map the evolution of chromosomes (Table 1). Here, in Table 1 we report that the predicted CT(s) with primary hierarchy is (are) largely syntenic to HSA19 (human chromosome 19, Group A’ – the CT with overall primary hierarchy), and/or HSA17 (human chromosome 17 – the CT with primary hierarchy in Group A’’, and secondary to HSA19). Most interestingly, in each of these studies, the hierarchical clustering of CTs delineates at least two primary clusters. The first of which has relatively high and diverse effective gene density (off-white to yellow block diagonal feature), and the second has a much lower but relatively uniform effective gene density (uniformly red block diagonal feature). Such a hierarchical pattern is similar to the high effective gene density block diagonal feature (Group A, Figure 1B), and the low effective gene density block diagonal feature (Group B, Figure 1B). As a further corroborative approach, the hierarchical position of human chrY (HSAY) is compared across the vertebrates: a recent ancestral primate: *Pan troglodytes* (Kuroki *et al.* 2006; Perry *et al.* 2007) to an evolutionarily distant ancestral vertebrate: *Gallus gallus* (Consortium 2004). The position of HSAY (Figure 1B), is juxtaposed between two large CT clusters, but the position of PTRY (the chrY in *Pan troglodytes)* is embedded in a CT sub-cluster, that is subordinate along with PTR2B (chr2B), PTR13 (chr13) and PTR18 (chr18) (Figure 5A), implying a significant difference in the inter-CT hierarchy. It is interesting that the orphan nature of chrY reported earlier for *Homo sapiens* (Figure 1A), is also embedded in the effective gene density matrix obtained from *Mus musculus* (Figure 5B), *Sus scrofa* (Figure 5C) and *Gallus gallus* (Figure 5D, with chromosome W) but definitely not in our more recent ancestor the *Pan troglodytes* (Figure 5A). Independent studies based on the genome sequence information have also reported that the HSAY significantly differs from PTRY, and also with the computed last common ancestral sequence (Hughes *et al.* 2010). Most interestingly, here we have identified this diversity between HSAY and PTRY without using nucleotide sequence information. Thus, the derived physical basis for a suprachromosomal ordering is applicable across diverse eukaryotic species, provided that their overall interphase CT arrangement is radial. We reiterate that our method succeeds in uncovering innate differences in chrY hierarchy vis-a-vis the rest of CTs in a genome as a system-level consequence, without invoking the sequence of chrY.

**Figure 5.**
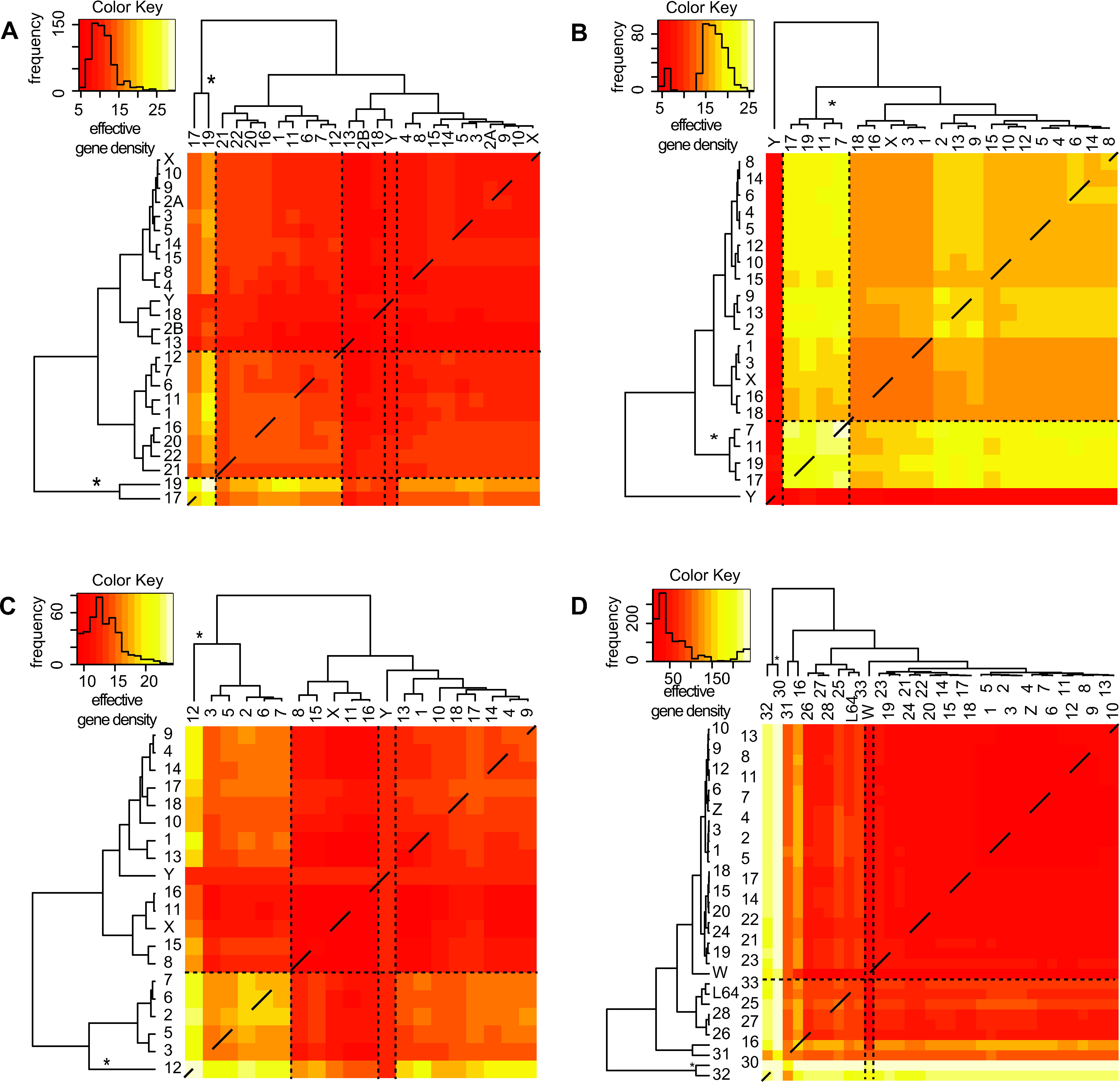
Heatmap and dendrogram obtained using effective gene density matrix from disparate vertebrate genomes. A heatmap and dendrogram represents the mathematical hierarchy of effective gene density matrix in diverse genomes that have radial chromosomal arrangement: (**A**) *Pan troglodytes*, (**B**) *Mus musculus*, (**C**) *Sus scrofa* and (**D**) *Gallus gallus*. Panel (**D**) represents the currently available annotated genes from NCBI Gene database and LGE64 is abbreviated as L64. The CT(s) with primary hierarchy in effective gene density space, and capturing the maximum diversity in inter-CT effective gene density having all colours: off-white (highest effective gene density) to red (least effective gene density) representing interior CT(s), and are marked (^*^). The chrY is delineated using a pair of dashed vertical lines in each instance. Inset histograms represent color keys.

### Gedanken experiments

In a thought experiment, consider the nuclear volume to be divided in five shells of nearly equal projected 2D sub-nuclear annular regions, as traditionally used in unbiased high-throughput imaging studies for the human nuclei (Boyle *et al.* 2001; Cremer *et al.* 2001; Bolzer *et al.* 2005; Mehta *et al.* 2013a; Mehta *et al.* 2013b). If we estimate the total number of ways we can progressively choose 46 CTs into those five shells, without considering any biological or stearic restrictions, then the total number of possible constellations is a tall order 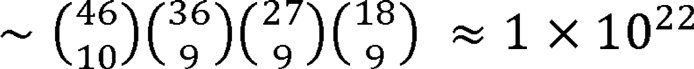. For this simple-minded calculation, we sought to accommodate nearly equal number of CTs in each of the five hypothetical concentric shells: 10 in one and 9 in each of the others. As all CTs do not occupy identical volumes, this is not an exact value but a theoretical exercise to demonstrate that a very large number of CT constellations may exist, substantiating a very plastic nature of inter-CT constellations.

Next, consider the reported probability distributions of individual CT. We estimate the fraction of the cell population, where each of the twenty-four distinct CTs is hypothetically localized at its most probable concentric shell. For simplicity, consider the marginal probability of individual CT occurrence as (similar magnitude for gene-rich CTs), then a marginal probability for all CTs in their most preferred shell is a very small percentage 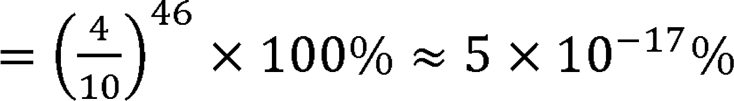. Hypothetically, it is plausible that the experimentally determined joint probabilities (not marginal probabilities) of select inter-CT pairs would be significantly larger compared to their marginal probabilities, our approximations suggests that there may be no unique constellation that can represent 1% from the same cell-type. However, we can rationalize that such highly likely constellations can well be more than times the second most populous constellation. Therefore, there may exist subgroups of similar constellations that predominate (Figure 3), and yet demonstrate the notion of a very high degree of plasticity in CT constellations, which is an important issue that has remained largely unaddressed by the community.

Now, we estimate the percentage of population that forty-five CTs occupying their most likely shell in various nuclei but with one of them occupying a least likely shell (say with hypothetical probability of 5%). The percentage of such nuclei will~ 6 × 10 ^−18^% of the entire population, which is an order of magnitude lower that the previous computation. Thus, a nontrivial fraction of nuclei may support CT constellations wherein one CT is in an unlikely location but all others occupy preferred positions (Figure 3A versus Figure 3B). Therefore, an empirical basis, which actually predates the full sequencing and annotation of the noncoding human genome, that suggests (i) “gene-rich” CTs, or (ii) small sized CTs occupy the nuclear interior, falls short in rationalizing a physical basis for a constellation of forty-six CTs. To further strengthen our hypothesis, we show the trend in the marginal probability of finding two leading gene-rich CTs, chr19 and chr17, in the interior versus the periphery. We do this by computing marginal probability of two and four of gene rich CTs and comparing those results obtained from two CTs that are localized to the nuclear periphery, chr5 and chr18 (Figure S2). Clearly, the marginal probabilities may report a significantly lower probability of simultaneous positioning of all gene rich CTs, or all gene poor CTs in the interior or the periphery.

### HSA19 dominates effective gene density hierarchy of protein-coding / noncoding genome

We wanted to contrast the exclusive inter-CT hierarchy from the protein-coding and noncoding genome. First, we re-derived effective (protein-coding) gene density matrix using the intrinsic parameters from the human protein-coding genome. Hence, the effective number of genes and effective gene density expressed in Equations 5, 6 and 8 at best represented 2% of the human genome. The heatmap representative of the inter-CT crosstalk for the coding genome is shown in Figure 6A. Although, the hierarchical clustering differs with that from the one in Figure 1, there are notable common features: (1) The chr19 has primary hierarchy, followed by chr17 exactly as in Figure 3, (2) The chrY is juxtaposed between two large subgroups of CTs (identical to Groups A and B in Figure 3). The differences between the dendrograms from Figure 1 and Figure 6A are: (1) All tertiary sub-clusters differ, (2) chr1 and chr12 are clustered with chr19, chr17, chr20 and chr22 (Figure 6A).

**Figure 6.**
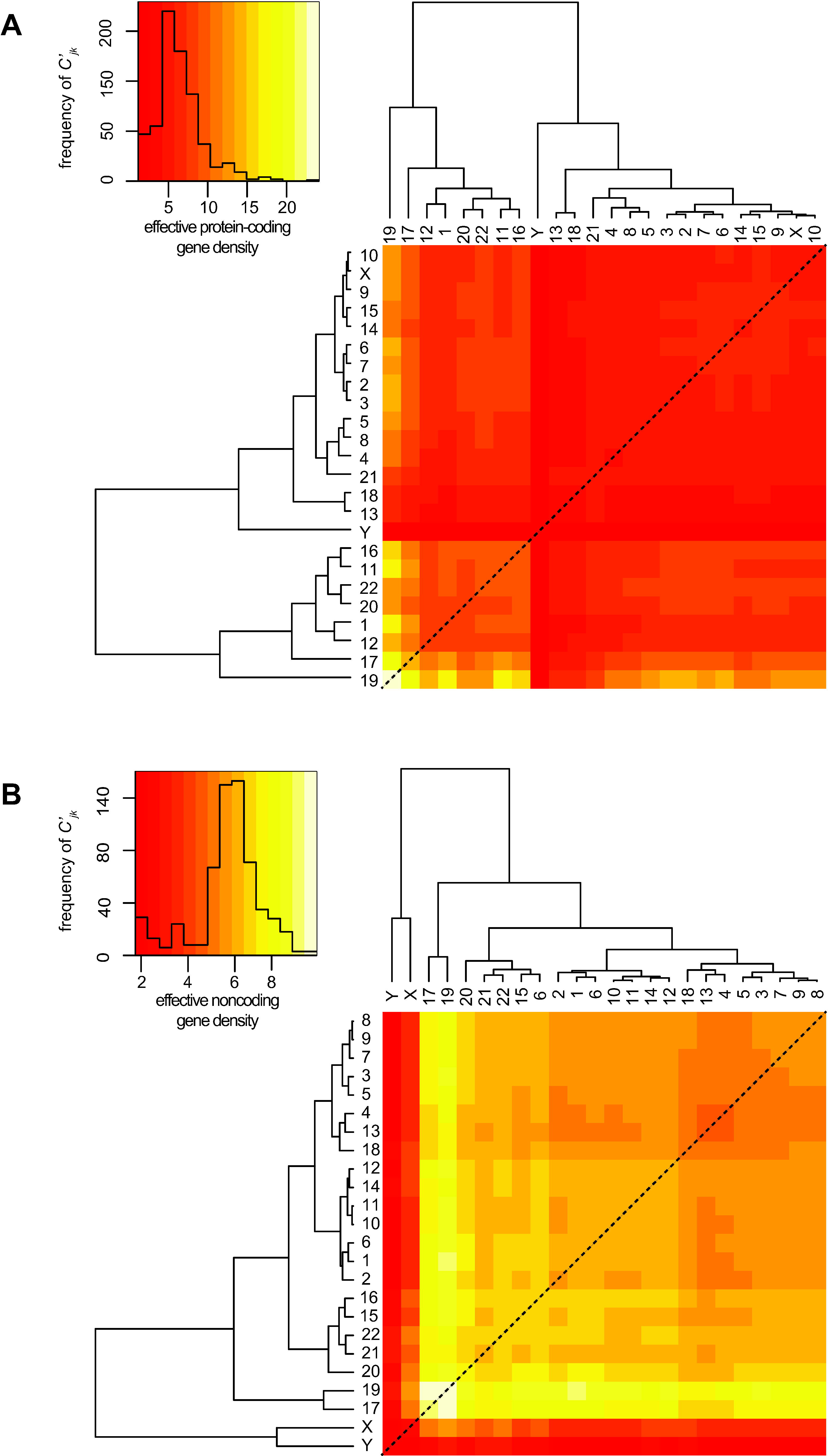
Hierarchical clustering of extrinsic inter-CT matrix derived from the coding / noncoding human genome. The hierarchy of extrinsic inter-CT effective gene density matrix as derived from the exclusive protein-coding genome (**A**), and the exclusive noncoding genome (**B**). Inset histograms represent color keys.

Following that, we computed an effective gene density matrix using the intrinsic parameters of the human noncoding genome. Its hierarchy (Figure 6B) in effective gene density is different to that from the coding genome (Figure 6A), or the full genome (Figure 1). The features differentiating dendrograms are: (1) The two most noncoding gene dense chromosomes: chr17 and chr19 cluster together, and the two sex chromosomes branch off from them, (2) The chrX and chrY cluster together, segregated from the autosomes (Figure 6B) and are most subordinate in the hierarchy - least effective noncoding gene density with all other CTs. Our overall results, suggest that within an effective gene density space, a differential mix of coding and noncoding parts of the genome might significantly alter effective gene density based inter-CT hierarchy.

### Saliency of hierarchy as obtained from effective gene density

In a given biological context, differential levels of activated and inactivated genes exist per chromosome, which may clearly influence both the intrinsic properties of a CT as well neighboring CTs via extrinsic couplings. During chromosomal translocations, the physical values of intrinsic parameters of affected CTs are also altered. An *in silico* analysis was implemented to estimate the incremental change in intrinsic parameters, which altered the original hierarchy and degeneracy in effective gene density. To assess the saliency and sensitivity in changing intrinsic parameters of this complex system, we only tuned one parameter at a time keeping others constant, generating hypothetical surrogate genomes. We computed the minimum decrement in the number of genes for each chromosome that led to an altered dendrogram in Figure 3. The change could be a reorganization of the hierarchy within at least one subgroup. The results of such sensitivity analysis for all chromosomes are presented in the Table S3. We observed that the decrement of genes required for a change in dendrogram for each chromosome was variable. In summary, a sensitivity analysis substantiates that primarily small changes (~1%) and few large changes (~10%) in the number of genes within a chromosome adversely affect the original hierarchy and degeneracy of CTs in effective gene density space. However, in any case, the decrement computed as a percent change in the genome was rather small (0.1–1.0%), which demonstrates that hierarchy dendrogram is a sensitive construct representative of the whole human genome organization, with regard to a CT constellation in the nucleus. Our saliency analysis also supports the inherent plasticity among constellations. Clearly, omission of a hundred genes is not feasible in a biological context, but this *in silico* experiment helps uncover the level of sensitivity of effective gene density hierarchy in a pan-nuclear setting.

## Discussion

At an empirical level, barring exceptions, intrinsic parameters coding gene density or/and chromosome size have been implicated to predict CT arrangement from FISH studies (Croft *et al.* 1999; Sun *et al.* 2000; Boyle *et al.* 2001; Cremer *et al.* 2001; Bolzer *et al.* 2005). It has also been reported that relatively small-sized CTs, independent of their protein-coding gene density, are localized to the nuclear interior, but larger ones are at the periphery (Sun *et al.* 2000; Cremer *et al.* 2001; Bolzer *et al.* 2005), again barring exceptions. As a specific example, inconsistent CT locations between two widely studied cell-types, fibroblasts and lymphocytes are considered (Table S1) (Boyle *et al.* 2001; Mehta *et al.* 2013b). Here, we instantiate how intrinsic parameters do not completely represent the physical basis of spatial CT arrangement in a densely packed milieu of human nuclei. (1) Spatial positions of chr21 and chrY, both small-size CTs with low gene density, are inconsistent in fibroblast and lymphocyte nuclei. Both CTs are largely at the nuclear interior in fibroblasts but also at the periphery in lymphocytes. (2) Inconsistency in the spatial locations of seven other CTs (chr1, chr5, chr6, chr8, chr15, chr16 and chr20) in fibroblasts versus lymphocytes. (3) Consistent (or weakly consistent) consensus locations obtained for chr10, chr11 and chr14 in these two cell-types. Interestingly, unlike most other CTs, these do not have a preferred spatial territory in the nuclei (interior versus intermediate versus periphery). (4) Most importantly, in the realm of protein-coding gene density or size-based segregation, the threshold of high versus low intrinsic parameter(s) that segregates interior versus peripheral CTs is qualitative or at best empirical, and lacks mathematical rigor because individual CT constellations have not been mapped. On the other hand, our theory supports an overall unique nonrandom hierarchical arrangement of CTs (a CT constellation), which arises along with permissible degeneracy and positional anisotropy, a result that has been experimentally confirmed using Hi-C techniques (Lieberman-Aiden *et al.* 2009; Kalhor *et al.* 2012; Nagano *et al.* 2013). Moreover, our theoretical formalism derived at the systems-level suprachromosomal scale, provides the mathematical basis for a self-organized CT arrangement due to nearest-neighbor CT effects within the nucleus, substantiating a previously proposed hypothesis (Misteli 2001; Misteli 2009).

Matrix algebra have been extensively used to describe systems-level information that may be innocuous at times, such as in molecular phylogenetics, involving sequence evolution as illustrated via phylogenetic trees (Felsenstein 2004). In an unrelated study, systems-level approach has computed differential gene expressions obtained from microarray data by deriving “eigengenes” and “eigenarrays” via vectors and matrices to identify coexpressed over-active and under-active regulatory genes in genome-wide studies (Alter *et al.* 2000). While in the cited study, matrix formalism has been used to decompose the expression data; a converse approach has been used here since we unify the intrinsic chromosomal parameters with one another to uncover the systems-level crosstalk among CTs.

We have demonstrated that the extrinsic inter-CT couplings (effective gene density) derived using our systems-level theory substantiates the hierarchy, degeneracy, and constrains spatial CT arrangement. We surmise that such biological hierarchy ensures a self-organization in human nuclei with forty-six CTs. In fact for a self-organizing nucleus with forty-six CTs, the multiplicity of inter-CT coupling emphasizes that such systems are dynamic and require coupling and decoupling among themselves to sustain radial arrangement. Moreover, there is compelling evidence from Hi-C studies to suggest cell-to-cell variations in topological conformation of CTs (Nagano *et al.* 2013), and that CTs may also have preferential orientation with respect to landmarks such as the nuclear envelop (Schmalter *et al.* 2014), supporting neighborhood influence (mathematically demonstrated here in our paired chromosome’s gene count formalism). Here, as a mathematical exercise, we have investigated three paradigms that involved (i) the whole genome, protein-coding and noncoding parts of the genome (Figure 1 and 4), (ii) protein-coding part of the genome exclusively (Figure 6A), and (iii) noncoding part of the genome exclusively (Figure 6B). All three paradigms support the hierarchy of extrinsic effective gene density matrix, which led us to unravel the physical principle of a selforganized assembly of CTs. We have approximated the inter-CT crosstalk by using the first-order interactions or nearest-neighbor CTs (Materials and Methods). Interestingly, chrY has the most subordinate hierarchy in each of these three paradigms, reinforcing that the physical basis for CT constellations at the suprachromosomal level is identical for the 46,XY and 46,XX diploid genome. Although we identified patterns of similarity and dissimilarity in the extrinsic inter-CT couplings for these three paradigms, we obtained results that support an overall hierarchy and plasticity of inter-CT constellations. The uniqueness in the results of the three paradigms discussed above suggests instances when the three paradigms may coexist to varying degrees as dictated by the biology of the nucleus, whose detailed analysis is beyond the scope of our current study. We reiterate that the abstract effective gene density space defined by the differential mix of coding versus noncoding human genome might significantly alter inter-CT coupling hierarchy. In theory, if one was to extend this model to incorporate three nearest-neighbor CT interactions, additional genomic constraints will be described, implying much tighter constraints among genomic parameters.

In conclusion, the effective gene density matrix of the human genome unifies intrinsic parameters of CTs and mathematically constraints their spatial arrangement. The anisotropy associated with effective gene density, in our *in vivo* representation, provides a physical basis for a non-random and self-organized radial CT arrangement in a densely packed nuclear milieu. We have corroborated and rationalized our findings with available experimental data, and substantiated our hypothesis in disparate mammalian species that extrinsic effective gene density matrix constrains spatial CT arrangement in interphase nuclei. We surmise that the effective gene density based anisotropy in the nuclear milieu largely dictates spatial hierarchy and associated plasticity of CT constellations. We also hypothesize that a high plasticity of CT constellations, within a given clonal population of cells, might set up cellular heterogeneity, as reported in single cell Hi-C studies (Nagano *et al.* 2013). However, the cell-type specific nuclear architectural components, via lamin and other nuclear-matrix mediated interactions, (van Steensel and Henikoff 2000; Pickersgill *et al.* 2006; Guelen *et al.* 2008) contribute towards the maintenance of CT arrangement across cell divisions. Finally, we also applied this systems-level theory across disparate mammalian species with annotated genomes, and identified the interior core CTs in their nuclei. Our results provide a novel perspective on the inter-CT arrangement (*in vivo* evolution of CT constellations in 3D) within vertebrate nuclei in the broader evolutionary context, analogous to the *in vitro* evolution of individual human chromosomes in a 1D framework. Therefore, an identical suprachromosomal physical basis, which constrains a self-organized spatial CT arrangement in human nuclei, is also conceptualized across disparate vertebrate species in the context of a radial CT arrangement.

## Acknowledgements

The authors thank Professor G Ravindrakumar (Department of Nuclear & Atomic Physics, TIFR) and Dr. Ravi Venkatramani (Department of Chemical Sciences, TIFR) for a critical reading of the manuscript and suggestions. SNF acknowledges Carson C. Chow (NIH/NIDDK/LBM) for his inputs, Peter Cooper (NIH/NLM/NCBI) and Wayne Matten (NIH/NLM) for help with access to annotated genes from NCBI’s Gene and Mapview databases. This work was supported by funding from: TIFR-Department of Atomic Energy (BJR, SNF), Grant 12P-0123 (BJR), and Sir JC Bose Award Fellowship, Department of Science and Technology, Government of India: Grant 10X-217 (BJR).

## Symbols and mathematical representations in vector space

In this report, 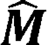denotes a matrix,❘***C***⟩ denotes a column vector, and ⟨***R***❘denotes a row vector, such that 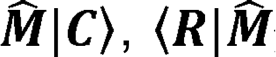, and ⟨***R***❘***C***⟩ denote inner products and ❘***C***⟩⟨***R***❘ denotes an outer product. The labels ***C*** and ***R*** are parameters (either intrinsic or extrinsic parameters) that represent those vectors. Notations used herein are similar to that in Alter O., Brown P.O. and Botstein D. (*Proc. Nat. Acad. Sci., USA 2000*).

## References

Alter,O., P. O. Brown and D. Botstein, 2000 Singular value decomposition for genome-wide expression data processing and modeling. Proc Natl Acad Sci U S A 97: 10101–10106.

Barbieri,M., M. Chotalia, J. Fraser, L.M. Lavitas, J. Dostie et al., 2012 Complexity of chromatin folding is captured by the strings and binders switch model. Proc Natl Acad Sci U S A 109: 16173–16178.

Bickmore,W. A., 2013 The spatial organization of the human genome. Annu Rev Genomics Hum Genet 14: 67–84.

Bickmore,W. A., and P. Teague, 2002 Influences of chromosome size, gene density and nuclear position on the frequency of constitutional translocations in the human population. Chromosome Res 10: 707–715.

Bickmore,W. A., and B. van Steensel, 2013 Genome architecture: domain organization of interphase chromosomes. Cell 152: 1270–1284.

Blackstone,T., R. Scharein, B. Borgo, R. Varela, Y. Diao et al., 2011 Modeling of chromosome intermingling by partially overlapping uniform random polygons. J Math Biol 62: 371–389.

Bolzer,A., G. Kreth, I. Solovei, D. Koehler, K. Saracoglu et al., 2005 Three-dimensional maps of all chromosomes in human male fibroblast nuclei and prometaphase rosettes. PLoS Biol 3: e157.

Boyle,S., S. Gilchrist, J. M. Bridger, N. L. Mahy, J. A. Ellis et al., 2001 The spatial organization of human chromosomes within the nuclei of normal and emerin-mutant cells. Hum Mol Genet 10: 211–219.

Branco,M. R., and A. Pombo, 2006 Intermingling of chromosome territories in interphase suggests role in translocations and transcription-dependent associations. PLoS Biol 4: e138.

Bridger,J. M., S. Boyle, I. R. Kill and W.A. Bickmore, 2000 Re-modelling of nuclear architecture in quiescent and senescent human fibroblasts. Curr Biol 10: 149152.

Cavalli,G., and T. Misteli, 2013 Functional implications of genome topology. Nat Struct Mol Biol 20: 290–299.

Consortium,I. C. G. S., 2004 Sequence and comparative analysis of the chicken genome provide unique perspectives on vertebrate evolution. Nature 432: 695716.

Cook,P. R., and D. Marenduzzo, 2009 Entropic organization of interphase chromosomes. J Cell Biol 186: 825–834.

Cortez,D., R. Marin, D. Toledo-Flores, L. Froidevaux, A. Liechti et al., 2014 Origins and functional evolution of Y chromosomes across mammals. Nature 508: 488–493.

Cremer,M., J. von Hase, T. Volm, A. Brero, G. Kreth et al., 2001 Non-random radial higher-order chromatin arrangements in nuclei of diploid human cells. Chromosome Res 9: 541–567.

Cremer,T., C. Cremer, T. Schneider, H. Baumann, L. Hens et al., 1982 Analysis of chromosome positions in the interphase nucleus of Chinese hamster cells by laser-UV-microirradiation experiments. Hum Genet 62: 201–209.

Cremer,T., and M. Cremer, 2010 Chromosome territories. Cold Spring Harb Perspect Biol 2: a003889.

Croft,J. A., J. M. Bridger, S. Boyle, P. Perry, P. Teague et al., 1999 Differences in the localization and morphology of chromosomes in the human nucleus. J Cell Biol 145: 1119–1131.

Dekker,J., and T. Misteli, 2015 Long-Range Chromatin Interactions. Cold Spring Harb Perspect Biol 7: a019356.

Dixon,J. R., S. Selvaraj, F. Yue, A. Kim, Y. Li et al., 2012 Topological domains in mammalian genomes identified by analysis of chromatin interactions. Nature 485: 376–380.

Dorier,J., and A. Stasiak, 2009 Topological origins of chromosomal territories. Nucleic Acids Res 37: 6316–6322.

Fatakia, S. N., I. S. Mehta and B. J. Rao, 2015 Unified theory of human genome reveals a constrained spatial chromosomal arrangement in interphase nuclei, pp. in arXiv:1509.08074 [q-bio.GN]. arXiv.org.

Federico,C., S. Saccone, L. Andreozzi, S. Motta, V. Russo et al., 2004 The pig genome: compositional analysis and identification of the gene-richest regions in chromosomes and nuclei. Gene 343: 245–251.

Felsenstein,J., 2004 Inferring Phylogenies. Macmillian Education.

Foster H. A., D. K. Griffin and J. M. Bridger, 2012 Interphase chromosome positioning in in vitro porcine cells and ex vivo porcine tissues. BMC Cell Biol 13: 30.

Fraser,J., C. Ferrai, A. M. Chiariello, M. Schueler, T. Rito et al., 2015 Hierarchical folding and reorganization of chromosomes are linked to transcriptional changes in cellular differentiation. Mol Syst Biol 11: 852.

Fudenberg,G., and L. A. Mirny, 2012 Higher-order chromatin structure: bridging physics and biology. Curr Opin Genet Dev 22: 115–124.

Ganai,N., S. Sengupta and G. I. Menon, 2014 Chromosome positioning from activity-based segregation. Nucleic Acids Res 42: 4145–4159.

Gerlich,D., J. Beaudouin, B. Kalbfuss, N. Daigle, R. Eils et al., 2003 Global chromosome positions are transmitted through mitosis in mammalian cells. Cell 112: 751–764.

Guelen,L., L. Pagie, E. Brasset, W. Meuleman, M. B. Faza et al., 2008 Domain organization of human chromosomes revealed by mapping of nuclear lamina interactions. Nature 453: 948–951.

Habermann,F. A., M. Cremer, J. Walter, G. Kreth, J. von Hase et al., 2001 Arrangements of macro- and microchromosomes in chicken cells. Chromosome Res 9: 569–584.

Halverson,J. D., J. Smrek, K. Kremer and A. Y. Grosberg, 2014 From a melt of rings to chromosome territories: the role of topological constraints in genome folding. Rep Prog Phys 77: 022601.

Heermann,D. W., H. Jerabek, L. Liu and Y. Li, 2012 A model for the 3D chromatin architecture of pro and eukaryotes. Methods 58: 307–314.

Hofmann,A., and D. W. Heermann, 2015 The role of loops on the order of eukaryotes and prokaryotes. FEBS Lett 589: 2958–2965.

Hughes,J. F., H. Skaletsky, T. Pyntikova, T. A. Graves, S. K. van Daalen et al., 2010 Chimpanzee and human Y chromosomes are remarkably divergent in structure and gene content. Nature 463: 536–539.

Kalhor,R., H. Tjong, N. Jayathilaka, F. Alber and L. Chen, 2012 Genome architectures revealed by tethered chromosome conformation capture and population-based modeling. Nat Biotechnol 30: 90–98.

Kreth,G., J. Finsterle, J. von Hase, M. Cremer and C. Cremer, 2004 Radial arrangement of chromosome territories in human cell nuclei: a computer model approach based on gene density indicates a probabilistic global positioning code. Biophys J 86: 2803–2812.

Kuroki,Y., A. Toyoda, H. Noguchi, T. D. Taylor, T. Itoh et al., 2006 Comparative analysis of chimpanzee and human Y chromosomes unveils complex evolutionary pathway. Nat Genet 38: 158–167.

Lanctot,C., T. Cheutin, M. Cremer, G. Cavalli and T. Cremer, 2007 Dynamic genome architecture in the nuclear space: regulation of gene expression in three dimensions. Nat Rev Genet 8: 104–115.

Lieberman- Aiden,E., N. L. van Berkum, L. Williams, M. Imakaev, T. Ragoczy et al., 2009 Comprehensive mapping of long-range interactions reveals folding principles of the human genome. Science 326: 289–293.

Marti-Renom,M. A., and L.A. Mirny, 2011 Bridging the resolution gap in structural modeling of 3D genome organization. PLoS Comput Biol 7: e1002125.

Mayer,R., A. Brero, J. von Hase, T. Schroeder, T. Cremer et al., 2005 Common themes and cell type specific variations of higher order chromatin arrangements in the mouse. BMC Cell Biol 6: 44.

Mehta,I., S. Chakraborty and B. J. Rao, 2013a IMACULAT – an open access package for the quantitative analysis of chromosome localization in the nucleus. PLoS One 8: e61386.

Mehta,I. S., M. Kulashreshtha, S. Chakraborty, U. Kolthur-Seetharam and B. J. Rao, 2013b Chromosome territories reposition during DNA damage-repair response. Genome Biol 14: R135.

Misteli,T., 2001 The concept of self-organization in cellular architecture. J Cell Biol 155: 181–185.

Misteli,T., 2009 Self-organization in the genome. Proc Natl Acad Sci U S A 106: 68856886.

Misteli,T., 2010 Higher-order genome organization in human disease. Cold Spring Harb Perspect Biol 2: a000794.

Munkel,C., R. Eils, S. Dietzel, D. Zink, C. Mehring et al., 1999 Compartmentalization of interphase chromosomes observed in simulation and experiment. J Mol Biol 285: 1053–1065.

Munkel,C., and L. Mirny, 1998 Chromosome structure predicted by a polymer model. Phys Rev E Stat Nonlin Soft Matter Phys 57: 5888.

Nagano,T., Y. Lubling, T. J. Stevens, S. Schoenfelder, E. Yaffe et al., 2013 Single-cell Hi-C reveals cell-to-cell variability in chromosome structure. Nature 502: 59–64.

NCBI, 2015 Database resources of the National Center for Biotechnology Information. Nucleic Acids Res 43: D6–17.

Nikiforova,M. N., J. R. Stringer, R. Blough, M. Medvedovic, J. A. Fagin et al., 2000 Proximity of chromosomal loci that participate in radiation-induced rearrangements in human cells. Science 290: 138–141.

Parada,L. A., P. G. McQueen and T. Misteli, 2004 Tissue-specific spatial organization of genomes. Genome Biol 5: R44.

Parada, L. A., P. G. McQueen, P. J. Munson and T. Misteli, 2002 Conservation of relative chromosome positioning in normal and cancer cells. Curr Biol 12: 16921697.

Perry,G. H., R. Y. Tito and B. C. Verrelli, 2007 The evolutionary history of human and chimpanzee Y-chromosome gene loss. Mol Biol E vol 24: 853–859.

Pickersgill,H., B. Kalverda, E. de Wit, W. Talhout, M. Fornerod et al., 2006 Characterization of the Drosophila melanogaster genome at the nuclear lamina. Nat Genet 38: 1005–1014.

Pombo,A., and N. Dillon, 2015 Three-dimensional genome architecture: players and mechanisms. Nat Rev Mol Cell Biol 16: 245–257.

Roix,J. J., P. G. McQueen, P.J. Munson, L. A. Parada and T. Misteli, 2003 Spatial proximity of translocation-prone gene loci in human lymphomas. Nat Genet 34: 287–291.

Rosa,A., and R. Everaers, 2008 Structure and dynamics of interphase chromosomes. PLoS Comput Biol 4: e1000153.

Schmalter,A. K., A. Kuzyk, C. H. Righolt, M. Neusser, O. K. Steinlein et al., 2014 Distinct nuclear orientation patterns for mouse chromosome 11 in normal B lymphocytes. BMC Cell Biol 15: 22.

Sengupta,K., J. Camps, P. Mathews, L. Barenboim-Stapleton, Q. T. Nguyen et al., 2008 Position of human chromosomes is conserved in mouse nuclei indicating a species-independent mechanism for maintaining genome organization. Chromosoma 117: 499–509.

Sexton,T., and G. Cavalli, 2015 The Role of Chromosome Domains in Shaping the Functional Genome. Cell 160: 1049–1059.

Smith,E. M., B. R. Lajoie, G. Jain and J. Dekker, 2016 Invariant TAD Boundaries Constrain Cell-Type-Specific Looping Interactions between Promoters and Distal Elements around the CFTR Locus. Am J Hum Genet 98: 185–201.

Sun,H. B., J. Shen and H. Yokota, 2000 Size-dependent positioning of human chromosomes in interphase nuclei. Biophys J 79: 184–190.

Tanabe,H., F. A. Habermann, I. Solovei, M. Cremer and T. Cremer,2002a Non-random radial arrangements of interphase chromosome territories: evolutionary considerations and functional implications. Mutat Res 504: 37–45.

Tanabe,H., S. Muller, M. Neusser, J. von Hase, E. Calcagno et al., 2002b Evolutionary conservation of chromosome territory arrangements in cell nuclei from higher primates. Proc Natl Acad Sci U S A 99: 4424–4429.

Tumbar,T., and A. S. Belmont, 2001 Interphase movements of a DNA chromosome region modulated by VP 16 transcriptional activator. Nat Cell Biol 3: 134–139.

van Steensel,B., and S. Henikoff, 2000 Identification of in vivo DNA targets of chromatin proteins using tethered dam methyltransferase. Nat Biotechnol 18: 424–428.

Vasquez,P. A., and K. Bloom, 2014 Polymer models of interphase chromosomes in Nucleus.

Vettorel,T., A. Y. Grosberg and K. Kremer, 2009 Statistics of polymer rings in the melt: a numerical simulation study. Phys Biol 6: 025013.

Walter,J., L. Schermelleh, M. Cremer, S. Tashiro and T. Cremer, 2003 Chromosome order in HeLa cells changes during mitosis and early G1, but is stably maintained during subsequent interphase stages. J Cell Biol 160: 685–697.

Waterston,R. H., K. Lindblad-Toh, E. Birney, J. Rogers, J. F. Abril et al., 2002 Initial sequencing and comparative analysis of the mouse genome. Nature 420: 520562.

Zhao,Z., G. Tavoosidana, M. Sjolinder, A. Gondor, P. Mariano et al., 2006 Circular chromosome conformation capture (4C) uncovers extensive networks of epigenetically regulated intra- and interchromosomal interactions. Nat Genet 38: 1341–1347.

Zink,D., T. Cremer, R. Saffrich, R. Fischer, M. F. Trendelenburg et al., 1998 Structure and dynamics of human interphase chromosome territories in vivo. Hum Genet 102: 241–251.

